# Discovery of new antibacterial accramycins from a genetic variant of the soil bacterium, *Streptomyces* sp. MA37

**DOI:** 10.1101/2020.09.14.295873

**Authors:** Fleurdeliz Maglangit, Yuting Zhang, Kwaku Kyeremeh, Hai Deng

## Abstract

Continued mining of natural products from the strain *Streptomyces* sp. MA37 in our laboratory led to the discovery of a minor specialised metabolite (SM) called accramycin A. Owing to its low yield (0.2mg/L) in the wild type strain, we investigated the roles of regulatory genes in the corresponding biosynthetic gene cluster (*acc* BGC) through gene inactivation with the aim of improving the titre of this compound. One of the resulting mutants (Δ*accJ*) dramatically upregulated the production of accramycin A **1** by 330-fold (66mg/L). Furthermore, ten new metabolites, accramycins B-K **2-11**, were discovered, together with two known compounds, naphthacemycin B_1_ **12** and fasamycin C **13** from the mutant extract. This suggested that *accJ*, annotated as Multiple Antibiotic Resistance Regulator (MarR), is a negative regulator gene in the accramycin biosynthesis. Compounds **1-13** inhibited the Gram-positive pathogens (*S. aureus, E. faecalis*) and clinical isolates, *E. faecium* (K59-68 and K60-39), and *S. haemolyticus* with minimal inhibitory concentration (MIC) values in the range of 1.5-12.5µg/mL. Remarkably, compounds **1-13** displayed superior activity against K60-39 (MIC = 3.1-6.3µg/mL) than ampicillin (MIC = 25µg/mL), and offer promising potential for the development of accramycin-based antibiotics that target multidrug-resistant *Enterococcus* clinical isolates. Our results highlight the importance of identifying the roles of regulatory genes in natural product discovery.

## Introduction

Naphthacemycins and congeners are a class of rare aromatic polyketides consisting of a partially reduced 1-phenyltetracene pentacyclic core [1–3]. This group of specialised metabolites (SM) display potent activity again various multidrug-resistant Gram-positive pathogens, such as methicillin-resistant *Staphylococcus aureus* (MRSA) and vancomycin-resistant *Enterococcus faecalis* (VRE) [4]. Typical examples in this class are the recently discovered naphthacemycin congeners, fasamycins [4–6], formicamycins [4], and streptovertimycins [7] (Figure 1). It has been shown that fasamycin A inhibits type II fatty acid synthases (FASII), which are essential for bacterial cell viability, with a potent inhibitory effect against FASII *in vitro* in a low IC_50_ value (50 µg/mL) [5]. Due to their potent antibacterial activities, fasamycins have attracted the attention of medicinal chemists for total synthesis to study the structure-activity relationship of the fasamycin scaffold [8–10].

**Figure 1.**
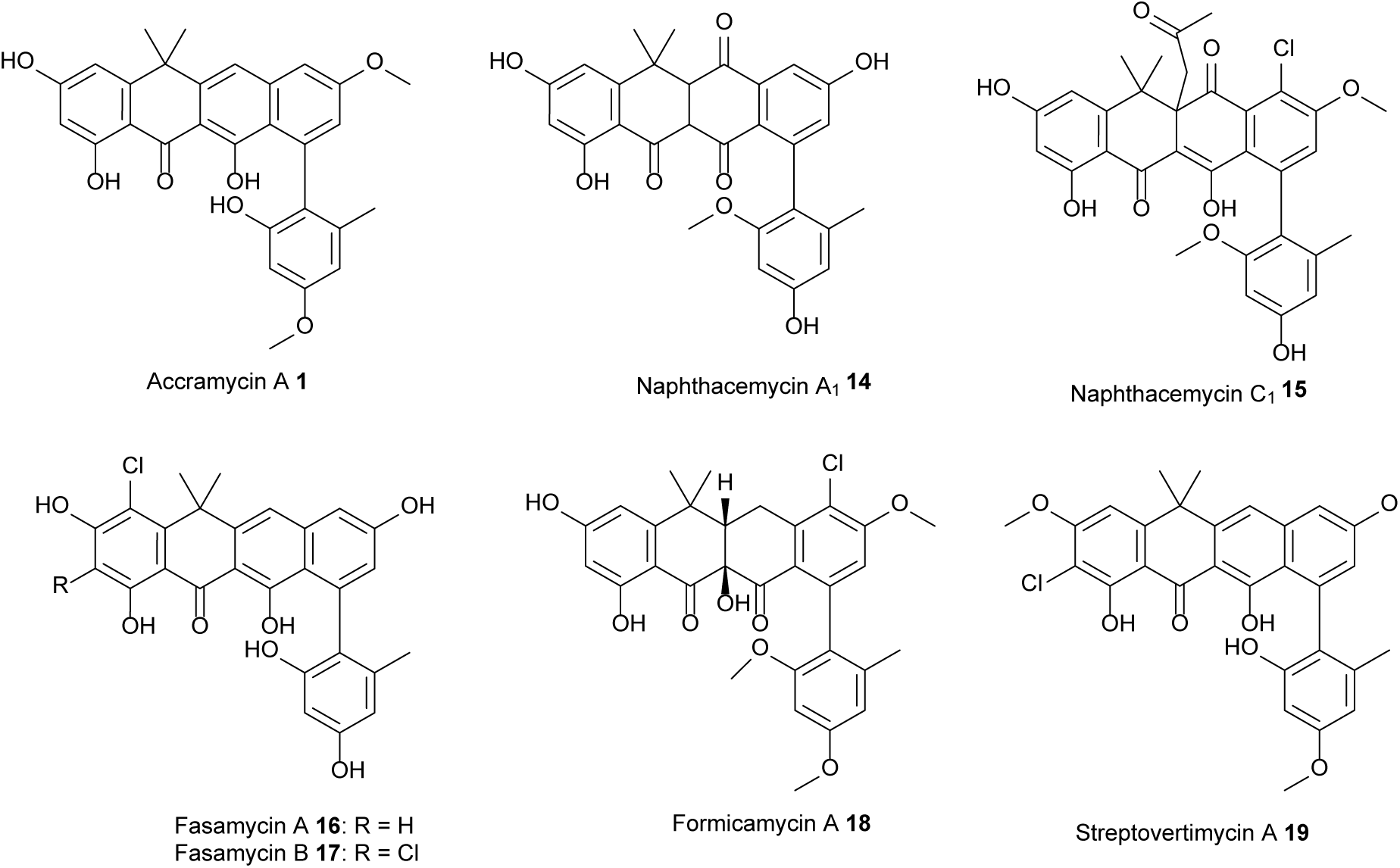
Structures of naphthacemycin-related specialised metabolites.

During our natural product screening programme, we found that the soil isolate, *Streptomyces* sp. MA37 is a highly prolific producer of SMs with great chemical diversity, including pyrrolizidine alkaloids [11], carbazoles [12–16], siderophores [17,18], and fluorinated compounds [19–24]. More recently, we discovered a new naphthacemycin congener, accramycin A **1**, as one of the minor SMs in the MA37 wild type (WT) strain (0.2 mg/L) [25]. Two mono-chlorinated accramycin derivatives were also tentatively assigned via molecular network analysis. Both derivatives, however, could not be isolated due to their minute quantities in the extract [25].

Based on the previous knowledge of fasamycins and formicamycins [4], the putative biosynthetic gene cluster (*acc* BGC) of **1** was identified [25]. The biosynthetic enzymes encoded in the *acc* BGC displays high amino acid (AA) similarity with the ones in fasamycins and formicamycins, including the FAD-dependent halogenation enzyme (AccV) which shares high AA identity to the chlorinase, ForV encoded in the fasamycin and formicamycin BGCs from *Streptomyces formicae* [4]. Unlike *S. formicae* that produces an array of mono-, di- and tri-chlorinated fasamycins and formicamycins, the MA37 WT strain appears to be less productive in terms of accramycin yield and chemical diversity, suggesting poor expression of the *acc* BGC in the MA37 WT strain [25]. It was hypothesized that this may be due to the presence of negative regulatory genes that suppresses the biosynthesis of accramycins.

Herein, we report the hyper accramycin producer of a MA37 variant by inactivating the putative regulatory gene (*accJ*) in the *acc* BGC, which encodes for Multiple Antibiotic Resistance Regulator (MarR). Subsequent chemical workup and structural elucidation allowed the discovery of more than ten new accramycin congeners, together with two known compounds, naphthacemycin B_1_ **12** [26] and fasamycin C **13** [4]. Except for compounds **3** and **4**, compounds **5-11** contain multi-chlorines installed at various positions of the accramycin scaffold, suggesting that the putative enzyme (AccV) is a promiscuous chlorinase. The major metabolite among these accramycin congeners is accramycin A with the estimated yield of 66 mg/L, a 330-fold increase over the one in the WT [25]. Compounds **1-13** inhibited the Gram-positive pathogens tested in this study with minimal inhibitory concentration (MIC) values in the range of 1.5-12.5µg/mL. Compounds **1-13** displayed superior activity against K60-39 (MIC = 3.1-6.3µg/mL) than ampicillin (MIC = 25µg/mL), and offer promising potential for the development of accramycin-based antibiotics that target multidrug-resistant *Enterococcus* clinical isolates.

## Results and Discussions

Bioinformatics analysis suggested that the *acc* BGC encodes four putative pathway-specific regulators, including LuxR (AccF), MarR (AccJ), LysR (AccI), and MerR (AccP) transcriptional regulators [25]. To assess their roles in the production of accramycins, we carried out the in-frame deletion of these four genes, generating four MA37 variants, respectively. The resulting mutants were then cultivated in ISP2 (7 days), followed by subsequent extraction to yield four crude extracts. Among the four crude extract samples, only the one from the Δ*acc*J variant displayed a significant metabolic profile in high-pressure liquid chromatography (HPLC) and high-resolution electrospray ionization mass spectrometry (HRESIMS) analyses compared with the WT (Figure S1). Of particular relevance is the presence of several new HPLC peaks with the characteristic UV pattern (226, 250, 286, 355, and 420nm), which have identical UV absorption maxima compared to the one of accramycin A we previously isolated [25].

To further confirm the identities of the newly emerged metabolites in Δ*accJ* variant, we set out a large-scale fermentation (2L) for chemical workup and structural elucidation. The crude extract was first fractionated through vacuum liquid chromatography to generate ten fractions. HPLC analysis of these ten fractions confirmed the presence of the accramycin constituents in fractions 3 (F3) and 4 (F4). The HPLC-UV targeted isolation approach afforded accramycin A **1** in a significantly improved titre (66mg/L), a 330-fold increase compared to the one from the MA37 WT (0.2mg/L) [25]. Additionally, several new accramycin analogues **2-11**, together with two known compounds naphthacemycin B_1_ **12** [26] and fasamycin C **13** [4] were isolated. Likewise, the titre of fasamycin C was enhanced in the mutant strain by 30-fold (26mg/L).

### Structure Elucidation

New accramycin analogues were obtained as yellowish to red powders. Inspection of the HR ESIMS data and MS/MS fragmentation pattern indicated that compounds **1-2** are non-halogenated, **3-4** have one chlorine atom, **5-7** have two chlorine atoms, **8-9** are trichloro-substituted, and **10-11** are tetrachlorinated. Thorough analysis of the UV, HR-ESIMS, and nuclear magnetic resonance (NMR) data of **1, 12** and **13**, indicated that compounds **1, 12** and **13** are known metabolites, accramycin A [25], naphthacemycin B_1_ produced by *Streptomyces* sp. KB-3346-5 (Figures S2-S21) [1,26], and fasamycin C produced by *S. formicae* (Fig. S20-25) [4], respectively. The structures of accramycins B-K **2-11** (Figure 2, Figures S22-S76) were elucidated by comparison of the observed UV, molecular formulae and NMR data with the reported data of **1, 12** and **13** (Tables S2-S3).

**Figure 2.**
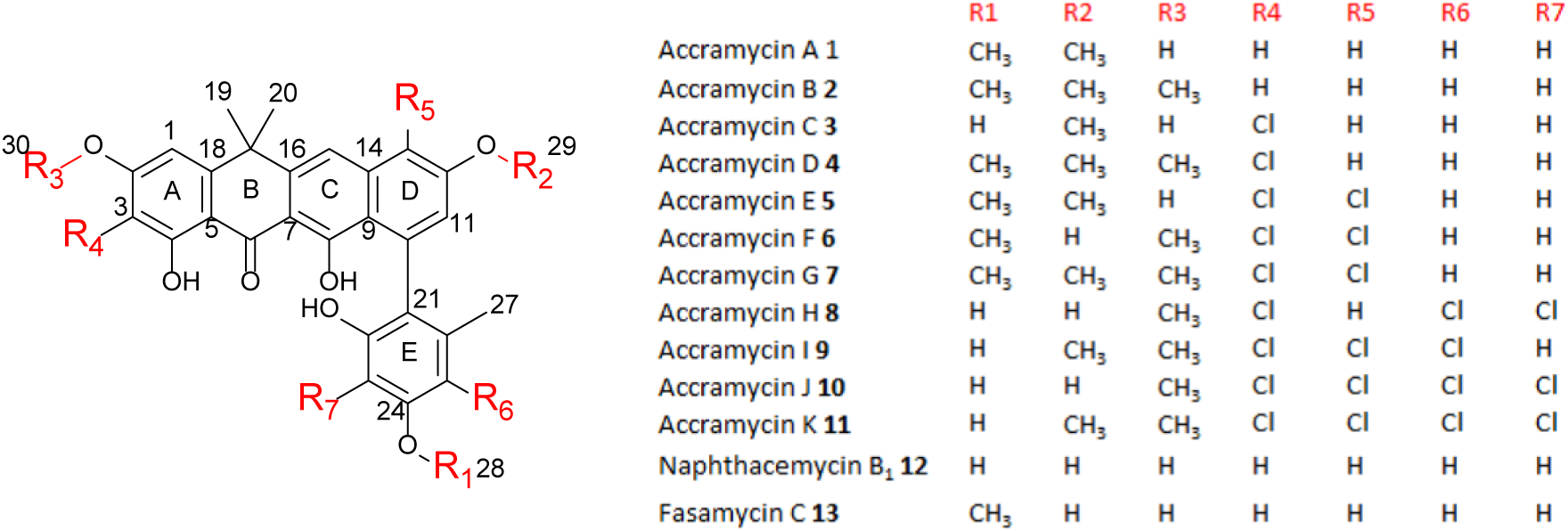
Accramycins isolated from *Streptomyces* sp. MA37 (Δ*accJ*) strain.

Compound **2** had the molecular formula C_30_H_28_O_7_ deduced by HRESIMS (observed [M+H]^+^ = 501.1898, *calcd* [M+H]^+^ = 501.1912 for C_30_H_29_O_7_^+^, Δ = -1.9ppm), suggesting 17 degrees of unsaturation. Detailed analysis of the LCMS, 1D and 2D NMR data of **2** revealed that it is similar to **1** [25] except for the presence of the methoxy moiety at C-2 in ring A, which was supported by the Heteronuclear Multiple Bond Correlation (HMBC) from H_3_-30 (δ_H_ 3.91) to C-2 (δ_C_ 166.5). Compound **2** was therefore identified as the new analogue, accramycin B.

The molecular formula C_28_H_23_ClO_7_ of compound **3** deduced by HRESIMS had 34 mass units more than **13**, which suggested that **3** was mono-chlorinated analogue of **13**. This was supported by the isotope pattern observed for **3** (*m/z* 507.1205 [M + H]^+^: 509.1166) in 3:1 ratio. The lower-field shifting of carbon signal (δ_C-3_ 106.6) in ring A indicated that the electronegative chlorine atom is attached to C-3 in the structure. This was further supported by the HMBC correlation from H-1 (δ_H_ 6.96) to C-3. The cross peak from H_3_-28 (δ_H_ 3.80) to C-24 (δ_C_ 159.0) established the connectivity of the methoxy functional group at C-24 in **3**.

HR ESIMS analysis of compound **4** deduced the molecular formula C_30_H_27_ClO_7_ (observed [M+H]^+^ = 535.1511, *calcd* [M+H]^+^ = 535.1518 for C_28_H_24_ClO_7_^+^, Δ = -1.4ppm), and indicated that **4** had 34 Da difference (Cl) from **2**. The additional chlorine atom was assigned at C-3 based on the HMBC correlation from H-1 (δ_H_ 6.97) to C-3 (δ_C_ 106.3).

Compounds E **5** and F **6** are isomers with the same molecular formula, C_29_H_24_Cl_2_O_7_ deduced by HR ESIMS. The 68 Da difference of **5-6** from **1** and the isotope fragmentation pattern (*m/z* 555.0978: 557.0944: 559.0925) in 9:6:1 ratio supported the presence of two chlorine atoms in the formula. Two chlorines were placed at C-3 and C-13 in **5-6**, based on the HMBC correlations from H-1 (δ_H_ 6.87) to C-3 (δ_C_ 106.8), H-11 (δ_H_ 7.05) to C-13 (δ_C_ 115.2), and H-1 (δ_H_ 7.00) to C-3 (δ_C_ 107.6), H-11 (δ_H_ 6.87) to C-13(δ_C_ 112.9), respectively. These compounds differed in the attachment of the two methoxy groups, where one was attached to C-24 in **5-6**, while the other was linked to C-12 in **5** and C-2 in **6**. This was confirmed by the correlations from H_3_-29 (δ_H_ 4.02) to C-12 (δ_C_ 155.8) and from H_3_-30 (δ_H_ 4.06) to C-2 (δ_C_ 161.5) in the HMBC spectra of **5** and **6**, respectively.

The ^1^H and ^13^C NMR spectra of compound **7** were similar to those of **6** except for ring D. The molecular formula of **7** C_30_H_26_Cl_2_O_7_ showed 14 mass units more than **6**, indicative of an additional methyl group in the structure which was assigned at C-12 based on the HMBC correlations from H_3_-29 (δ_H_ 4.02) to C-12 (δ_C_ 156.6).

Compound **8** is a tri-chlorinated isomer with the molecular formula, C_28_H_21_Cl_3_O_7_ deduced by HRESIMS. The H-NMR spectra of **8** revealed only one methoxy group which is assigned at C-2 based on the HMBC cross peaks between H_3_-30 (δ_H_ 4.05) and C-2 (δ_C_ 161.3). The three chlorines were designated based on the downfield carbon shifts observed in C-3 (δ_C_ 108.8), C-23 (δ_C_ 107.5) and C-25 (δ_C_ 113.2) in comparison with non-chlorinated **12**. The attachment of which was further supported by HMBC correlations from H-1 (δ_H_ 6.97) to C-3, H_3_-27 (δ_H_ 1.93) to C-23, and H-27 to C-25.

HRESIMS analysis established the molecular formula of compound **9**, C_29_H_23_Cl_3_O_7_ (*m/z* 589.0574 [M+H]^+^, 589.0582 *calcd* for C_29_H_24_Cl_3_O_7_^+^), and had 14 Da difference from **8**. The H-NMR spectra of **9** in comparison with **8** confirmed the presence of an additional methoxy substituent, and its location was established at C-12 of ring D based on the Nuclear Overhauser Enhancement Spectroscopy (NOESY) correlation between H_3_-29 (δ_H_ 4.02) and H-11 (δ_H_ 7.06). The three chlorines identified in the formula were assigned at C-3, C-13, and C-25 based on the HMBC correlation from H-1 (δ_H_ 7.00) to C-3 (δ_C_ 108.1), H-11 (δ_H_ 7.06) to C-13 (δ_C_ 115.5), and H_3_-27 (δ_H_ 1.98) to C-25 (δ_C_ 112.1 The ^1^H and ^13^C NMR spectra of compound **10** were similar to **8** except for ring D. HRESIMS analysis of **10** showed 34 mass units more than **8** indicating an additional chlorine substituent in the structure, which was assigned at C-13 based on the HMBC correlation from H-11 (δ_H_ 6.87) to C-13 (δ_C_ 113.8).

The ^1^H-NMR spectra of compound **11** showed an additional singlet signal in the methoxy region compared to **10**. The HMBC cross peak between H_3_-29 (δ_H_ 4.02) and C-12 (δ_C_ 156.2) designated the methoxy group at C-12 of ring D in the structure.

On the basis of the evidence in this study, compounds **2**-**11** were confirmed as new members of accramycin polyketides for which the names accramycin B-K are proposed, respectively.

### Biological Activity

Compounds **1-13** exhibited good activity against the Gram-positive pathogens, *Staphylococcus aureus* (ATCC 25923), *Enterococcus faecalis* (ATCC 29212), and clinical isolates *Enterococcus faecium* K59-68, *Enterococcus faecium* K60-39, and *Staphylococcus haemolyticus* 8-7A with minimum inhibitory concentration (MIC) values of 1.5-12.5 µg/mL. The presence of multiple chlorines at rings A, D, or E did not enhance the activity as observed in accramycins C-K consistent with previous findings [7]. The *O*-methyl bearing accramycin B **2** exhibited slightly higher MIC against all the tested pathogens. Conversely, the presence of free hydroxyl groups at C-12 and C-24 favours antibacterial activity as observed in accramycin J **10**. Compound **10** in comparison with naphthacemycin B_1_ **12** supported that the *O*-methyl at C-2 is a preferred structural feature for the activity. Furthermore, accramycin J **10** (MIC = 6.3 µg/mL) displayed 2-fold inhibitory activity over ampicillin (MIC = 12.5µg/mL) against the clinical isolate, *S. haemolyticus*. Notably, compounds **1-13** had superior activity (MIC = 3.1-6.3µg/mL) against *E. faecium* K60-39 than ampicillin (MIC = 25µg/mL). Thus, the accramycins represent potential therapeutic lead molecules for the development of potent drugs against ampicillin-resistant *Enterococcus* clinical strains. None of the compounds **1-13** displayed activity against the Gram-negative pathogens, *Escherichia coli* (ATCC 25922), and *Pseudomonas aeruginosa* (ATCC 27853), and the fungal pathogen *Candida albicans* (ATCC 10231) at the highest concentration tested (50µg/mL).

**Table 1.**
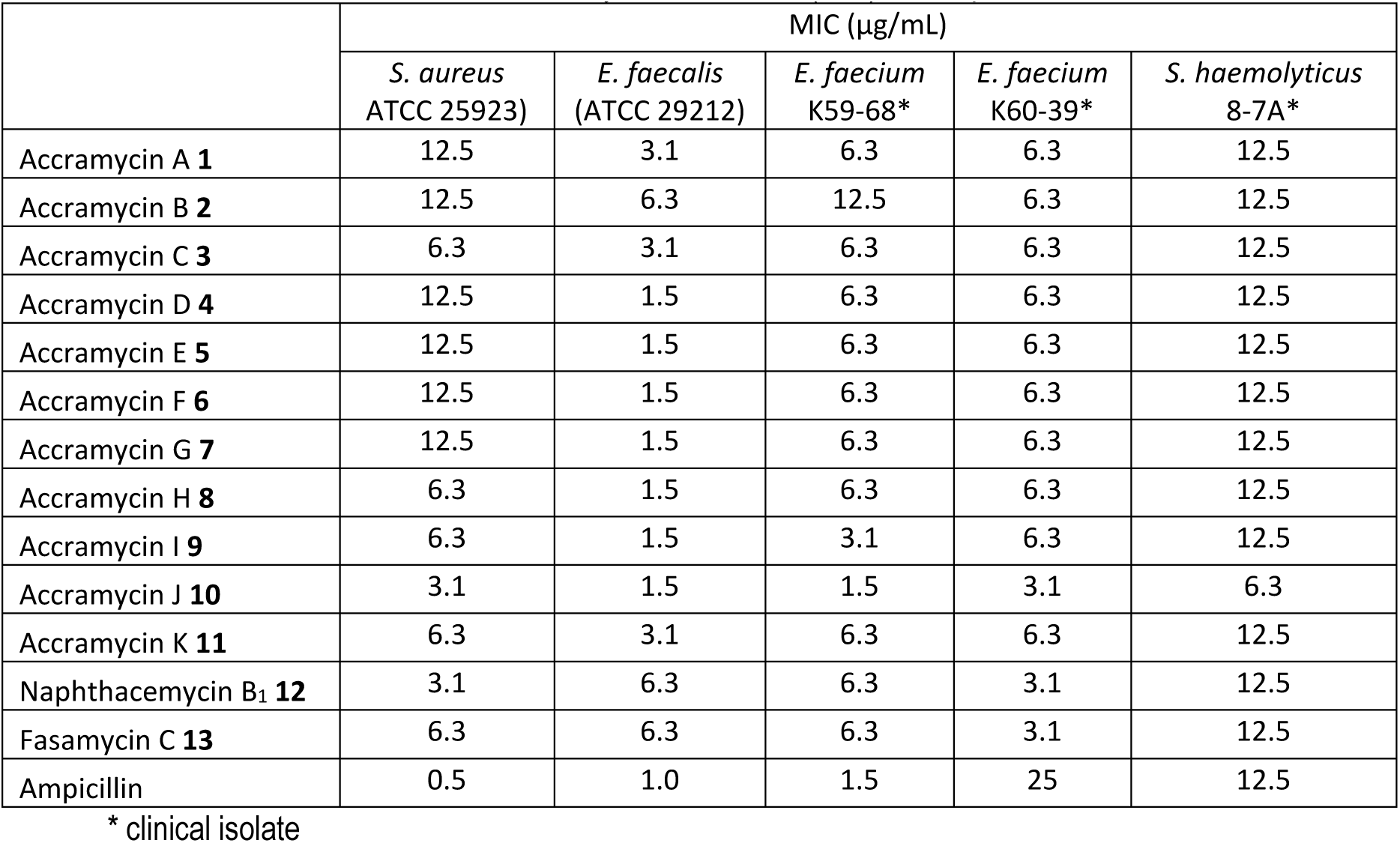
Minimum Inhibitory Concentration (MIC) of compounds **1-13**

|  | MIC (µg/mL) |  |  |  |  |
| --- | --- | --- | --- | --- | --- |
|  | *S. aureus*<br>ATCC 25923) | *E. faecalis*<br>(ATCC 29212) | *E. faecium*<br>K59-68* | *E. faecium*<br>K60-39* | *S. haemolyticus*<br>8-7A* |
| Accramycin A **1** | 12.5 | 3.1 | 6.3 | 6.3 | 12.5 |
| Accramycin B **2** | 12.5 | 6.3 | 12.5 | 6.3 | 12.5 |
| Accramycin C **3** | 6.3 | 3.1 | 6.3 | 6.3 | 12.5 |
| Accramycin D **4** | 12.5 | 1.5 | 6.3 | 6.3 | 12.5 |
| Accramycin E **5** | 12.5 | 1.5 | 6.3 | 6.3 | 12.5 |
| Accramycin F **6** | 12.5 | 1.5 | 6.3 | 6.3 | 12.5 |
| Accramycin G **7** | 12.5 | 1.5 | 6.3 | 6.3 | 12.5 |
| Accramycin H **8** | 6.3 | 1.5 | 6.3 | 6.3 | 12.5 |
| Accramycin I **9** | 6.3 | 1.5 | 3.1 | 6.3 | 12.5 |
| Accramycin J **10** | 3.1 | 1.5 | 1.5 | 3.1 | 6.3 |
| Accramycin K **11** | 6.3 | 3.1 | 6.3 | 6.3 | 12.5 |
| Naphthacemycin B<sub>1</sub> **12** | 3.1 | 6.3 | 6.3 | 3.1 | 12.5 |
| Fasamycin C **13** | 6.3 | 6.3 | 6.3 | 3.1 | 12.5 |
| Ampicillin | 0.5 | 1.0 | 1.5 | 25 | 12.5 |

\* clinical isolate

## Conclusions

In conclusion, inactivation of the *accJ* gene resulted in the production of thirteen naphthacemycin-type specialised metabolites in *Streptomyces* sp. MA37 *ΔaccJ* mutant. The structures of compounds **1-13** were deduced by HR-ESIMS and 1D and 2D NMR, including accramycins A-K **1-11** along with two known compounds, naphthacemycin B_1_ **12** and fasamycin C **13**. The fermentation titres of the isolated metabolites were significantly improved compared to the one from the wild type strain, particularly accramycin A **1** with estimated yield from 0.2mg/L (WT strain) to 66mg/L (Δ*accJ* mutant strain). This suggested that *accJ* is a repressor gene in the accramycin biosynthesis. Compounds **1-13** exhibited good activity against the Gram-positive pathogens, *S. aureus* and *E. faecalis*, and clinical isolates, *E. faecium* K59-68, *E. faecium* K60-39, and *S. haemolyticus* (MIC = 1.5-12.5µg/mL). Remarkably, compounds **1-13** displayed better inhibitory activity over ampicillin against K60-39 clinical isolate. Hence, the accramycin pharmacophore represents potential lead molecules for the development of potent antibiotics that target *Enterococcus* isolates.

## Materials and Methods

### General Experimental Procedures

Agilent 1260 Infinity was used for HPLC separation. IR spectra were obtained using a PerkinElmer Spectrum version 10.4.00 Fourier transform infrared (FTIR) spectrometer (2013) (UK) equipped with an Attenuated Total Reflection (ATR) diamond cell. HR-ESIMS were determined on LC-MS Thermal Science Mass Spectrometry (LTQ Orbitrap) coupled to a thermal instrument HPLC (Accela PDA detector, Accela PDA autosampler and Accela Pump, C18 Sunfire 150×46 mm waters). The NMR spectra were recorded on a Bruker AVANCE III HD 400 MHz (Ascend™ 9.4 Tesla) and/or Bruker AVANCE III HD 600 MHz (Ascend™ 14.1 Tesla) with Prodigy TCI™ cryoprobe.

### Strain, Genomic DNA, and media

The *Streptomyces* sp. MA37 strain was isolated from the soil sample collected from Legon, Ghana in West Africa [20]. Genomic DNA from MA37 was extracted and was used as PCR template for amplification of the up- and downstream fragments for the gene deletion. All media / broth for fermentation, solvents, and chemicals were obtained from Fisher Scientific (UK), unless otherwise stated.

### Construction of knockout vector

The left and right arms for the in-frame in *Streptomyces* sp. MA37 was generated by PCR using inverse primers (Table S1), with 15 bp overlap at their 5’ ends. The PCR mixture (25µL) contained genomic DNA of *Streptomyces* sp. MA37 (50ng) as the template, primers (1.0 µM), dNTP (0.1 mM, Novagen®), MgSO_4_ (0.1 mM), DMSO (4%), KOD buffer (10x), and KOD hot start DNA polymerase enzyme (0.5 µL). Reaction conditions consisted of an initial denaturation step at 95°C for 5 min followed by 30 cycles at 95°C for 30s, annealing at 60°C for 30s and a final extension step at 70°C for 2 minutes. The PCR fragments were verified on agarose gel.

The *E. coli* strain harbouring the pKC1139 vector was cultured overnight (37°C) in LB medium supplemented with kanamycin (50 µg/mL). The plasmid was extracted using QuickProtocol™ GeneJET™ plasmid miniprep kit, and linearized using the restriction enzymes, HindIII and EcoRI.

The two PCR fragments / inserts and the linearized vector were ligated by infusion cloning. The infusion reaction mixture (10µL) consisted of the purified PCR fragment (50-100ng), linearized vector (pKC1139) (50-100ng), 5x infusion HD enzyme premix (2µL), and water. The reaction was incubated (50°C, 15min). At the end of the reaction, the mixture was chilled on ice, followed by heat shock transformation into Stellar competent cells. Screening for the right construct was carried out by PCR using a pair of primers (Table S1).

### Conjugations between E. coli and Streptomyces sp. MA37

MA37 was cultured in YEME (yeast extract 3g, tryptone 5g, malt extract VWR 3g, glucose 10g, sucrose 103g in 1.0L milli-Q water) supplemented with glycine (0.5%) and magnesium chloride (10.0 mM) for 3 days. The mycelium was collected by centrifugation (3,000 × g, 10 min). The pellets were washed twice with sucrose (10.3%). Finally, the mycelium was resuspended in sucrose (10.3%, 5.0 mL).

The mycelium (0.5 mL) was mixed with *E. coli* S17-1 (0.5 mL), followed by centrifugation (5min). The supernatant was discarded, and the remaining mixture was plated onto SFM (soya flour mannitol) containing MgCl_2_ (10 mM). The plate was incubated at 28°C. After 16–20h, the plate was overlaid with water containing nalidixic acid (25µg/mL, 1mL) and apramycin (50µg/mL). Incubation was continued until the ex-conjugates appear (28°C, 3-5days).

### Screening for the Mutant Strain

The ex-conjugates were inoculated onto ISP2 (glucose 4g, yeast extract 4g, malt extract 10g, agar 20g, in 1L H_2_O) plates with apramycin (50µg/mL) and incubated at 37°C (2-3 generations). Using the streak plate method, each single colony was inoculated separately onto ISP2 and ISP2 with apramycin (50µg/mL). The colony that didn’t grow in the plate supplemented with apramycin was cultured in YEME for 3 days. The genomic DNAs of the ex-conjugates were extracted, followed by PCR verification using internal primers.

### Metabolic Profile of Mutant and WT strains

The seed cultures of variant strains were prepared by inoculating the glycerol stocks (10µL) in YEME (10mL), and incubating for 3 days (28°C, 180 rpm, Incu-shake FL16-2). The seed cultures were used to prepare small-scale cultures (50-mL) in 250-mL Erlenmeyer flasks (Pyrex(tm) borosilicate glass narrow neck flask containing ISP2 broth.

The cultures were incubated (7 days, 28°C, 180rpm), after which Diaion^®^ HP-20 (3g/50 mL solution) was added. Incubation was continued for the next 18-24h (28°C, 180 rpm). The resin was filtered, extracted exhaustively with methanol, followed by concentration using a rotary evaporator (Buchi Rotavapor R200, UK). Likewise, the MA37 WT strain was fermented and extracted as in the mutant strains. The mutants and WT extracts were subjected to mass spectrometric and HPLC-UV analyses monitored at λ450nm, and the chemical profiles of each were compared. Several peaks were observed in the *ΔaccJ* extract with the characteristic accramycin UV pattern (226, 250, 286, 355, and 420nm) not detected in the WT or other mutant strains.

### Fermentation, Extraction, Fractionation

Two-litre fermentation culture of the *ΔaccJ* mutant strain was carried out in ISP2 broth. Four 2.0-L baffled flasks (Corning™ polycarbonate) each containing 500mL ISP2, were inoculated with seed culture (1:100), and plugged with foam stoppers (Fisherbrand™ polyurethane). Fermentation, incubation and extraction of the *ΔaccJ* cultures were carried out as described above. The methanol extracts were combined, evaporated to dryness under reduced pressure to yield crude extract (7g), which was subjected to HR ESIMS analysis.

The crude extract was fractionated by vacuum liquid chromatography on silica gel 60 (Acros Organics™ultra-pure 60A 40-63u) eluting with a gradient of *n*-hexane−ethyl acetate-MeOH to give 10 subfractions (F1-F10). HPLC-UV analysis was carried out in all the fractions to screen and target the accramycin scaffold using semi-prep reversed-phase HPLC (ACE C18-HL 10µM 10 ×250 mm) equipped with a diode array detector (DAD) with spectral scanning between 200–550 nm. HPLC separation was carried out by solvent gradient method over a period of 45 min (flow rate 1.5mL/min, injection volume 10µL, solvent A - 95% H_2_O, 5% methanol, and 0.1% trifluoroacetic acid; solvent B - 100% methanol). The compounds of interest with the characteristic accramycin UV maxima (226, 250, 286, 355, and 420 nm) were observed in fractions F3-F4.

### Semi-prep HPLC Isolation

Further purification of F3-F4 fractions was carried out using semi-prep reversed-phase HPLC (C-18 ACE 10 µM 10 × 250 mm column) as described above, eluting with a 45 min gradient of 40-100% methanol. The purification afforded 11 accramycins A-K **1-11** along with two other known compounds, naphthacemycin B_1_ **12** [26], and fasamycin C **13** [4].

Accramycin A **1**: yield 131.4mg; deep yellow powder; UV (PDA) λ_max_: 225, 245, 290, 355, 420 nm; IR (neat) *v*_max_ (cm^−1^): 3350, 2946, 2834, 1681, 1607, 1284, 1202, 1026, 584; ^1^H, ^13^C NMR data, see Table S3; Molecular formula: C_29_H_26_O_7_ ; HRESIMS (positive mode) *m/z* calculated for C_29_H_27_O_7_^+^ [M + H]^+^ = 487.1751; observed [M + H]^+^ = 487.1748; Δ = -1.78 ppm
Accramycin B **2**: yield 6.12mg; deep yellow powder; UV (PDA) λ_max_: 250, 290, 315, 335, 350, 425 nm; IR (neat) *v*_max_ (cm^−1^): 3350, 2946, 2834, 1681, 1607, 1284, 1202, 1026, 584 ; Molecular formula: C_30_H_28_O_7_ ; ^1^H, ^13^C NMR data, see Table S3; HRESIMS (positive mode) *m/z* calculated for C_30_H_29_O_7_^+^ [M + H]^+^ = 501.1912; observed [M + H]^+^ = 501.1898; Δ = -1.978 ppm
Accramycin C **3**: yield 6.04mg; deep yellow powder; UV (PDA) λ_max_: 250, 280, 300, 355, 430nm; IR (neat) *v*_max_ (cm^−1^): 3362, 2922, 2848, 1679, 1612, 1443, 1203, 1149, 726 ; Molecular formula: C_28_H_23_ClO_7_ ; ^1^H, ^13^C NMR data, see Table S3; HRESIMS (positive mode) *m/z* calculated for C_28_H_24_ClO_7_^+^ [M + H]^+^ = 507.1205; observed [M + H]^+^ = 507.1207; Δ = 0.37 ppm.
Accramycin D **4**: yield 6.42mg; deep yellow powder; UV (PDA) λ_max_: 250, 290, 315, 335, 350, 425nm; IR (neat) *v*_max_ (cm^−1^) 3337, 2947, 2834, 1681, 1450, 1025, 634; ^1^H, ^13^C NMR data, see Table S3; Molecular formula: C_30_H_27_ClO_7_ ; HRESIMS (positive mode) *m/z* calculated for C_30_H_28_ClO_7_^+^ [M + H]^+^ = 535.1518; observed [M + H]^+^ = 535.1511; Δ = -1.8594ppm.
Accramycin E **5**: yield 6.68mg; deep yellow powder; UV (PDA) λ_max_: 250, 290, 315, 335, 350, 425 nm; IR (neat) *v*_max_ (cm^−1^) 3337, 2947, 2834, 1681, 1450, 1025, 634; ^1^H, ^13^C NMR data, see Table S3; Molecular formula: C_29_H_24_Cl_2_O_7_ ; HRESIMS (positive mode) *m/z* calculated for C_29_H_25_Cl_2_O_7_^+^ [M + H]^+^ = 555.0972; observed [M + H]^+^ = 555.0978; Δ = 1.18 ppm.
Accramycin F **6**: yield 7.04mg; deep yellow powder; UV (PDA) λ_max_: 250, 290, 315, 335, 350, 425 nm; IR (neat) *v*_max_ (cm^−1^) 3325, 2943, 2833, 1678, 1449, 1119, 1023, 635; ^1^H, ^13^C NMR data, see Table S3; Molecular formula: C_29_H_24_Cl_2_O_7_ ; HRESIMS (positive mode) *m/z* calculated for C_29_H_25_Cl_2_O_7_^+^ [M + H]^+^ = 555.0972; observed [M + H]^+^ = 555.0975; Δ = 0.63ppm.
Accramycin G **7**: yield 35.24mg; deep yellow powder; UV (PDA) λ_max_: 225, 250, 295, 320, 340, 350, 425 nm; IR (neat) *v*_max_ (cm^−1^) 3399, 2917, 2851, 1688, 1606, 1423, 1322, 1205; ^1^H, ^13^C NMR data, see Table S3; Molecular formula: C_30_H_26_Cl_2_O_7_ ; HRESIMS (positive mode) *m/z* calculated for C_30_H_27_Cl_2_O_7_^+^ [M + H]^+^ = 569.1128; observed [M + H]^+^ = 569.1127; Δ = - 0.29 ppm.
Accramycin H **8**: yield 6.24mg; deep yellow powder; UV (PDA) λ_max_: 250, 290, 310, 355, 425 nm; IR (neat) *v*_max_ (cm^−1^) 3337, 2946, 1678, 1448, 1204, 1021, 644; ^1^H, ^13^C NMR data, see Table S3; Molecular formula: C_28_H_21_Cl_3_O_7_ ; HRESIMS (positive mode) *m/z* calculated for C_28_H_22_Cl_3_O_7_^+^ [M + H]^+^ = 575.0426; observed [M + H]^+^ = 575.0420; Δ = -0.99ppm.
Accramycin I **9**: yield 13.58mg; deep yellow powder; UV (PDA) λ_max_: 250, 290, 315, 340, 350, 425 nm; IR (neat) *v*_max_ (cm^−1^) 3399, 2921, 2849, 1680, 1442, 1196, 1139; ^1^H, ^13^C NMR data, see Table S5; Molecular formula: C_29_H_23_Cl_3_O_7_ ; HRESIMS (positive mode) *m/z* calculated for C_29_H_24_Cl_3_O _7_^+^ [M + H]^+^ = 589.0582; observed [M + H]^+^ = 589.0574; Δ = -1.3217 ppm.
Accramycin J **10**: yield 15.04mg; deep yellow powder; UV (PDA) λ_max_: 250, 290, 315, 340, 350, 425 nm; IR (neat) *v*_max_ (cm^−1^) 3427, 1688, 1601, 1439, 1328, 1204, 1139; ^1^H, ^13^C NMR data, see Table S3; Molecular formula: C_28_H_20_Cl_4_O_7_ ; HRESIMS (positive mode) *m/z* calculated for C_28_H_21_Cl_4_O_7_^+^ [M + H]^+^ = 609.0036; observed [M + H]^+^ = 609.0040; Δ = 0.722 ppm.
Accramycin K **11**: yield 6.84mg; deep yellow powder; UV (PDA) λ_max_: 225, 250, 295, 320, 340, 350, 425 nm; IR (neat) *v*_max_ (cm^−1^) 3427, 1688, 1601, 1439, 1328, 1204, 1139 ; ^1^H, ^13^C NMR data, see Table S3; Molecular formula: C_29_H_22_Cl_4_O_7_ ; HRESIMS (positive mode) *m/z* calculated for C_29_H_23_Cl_4_O_7_^+^ [M + H]^+^ = 623.0192; observed [M + H]^+^ = 623.0184; Δ = 0.405 ppm.

Naphthacemycin B_1_ **12**: yield 86.38mg; reddish powder; UV (PDA) λ_max_: 245, 290, 355, 420 nm; IR (neat) *v*_max_ (cm^−1^) 3337, 2946, 1678, 1448, 1204, 1021, 644; ^1^H, ^13^C NMR data, see Table S3; Molecular formula: C_27_H_22_O_7_ ; HRESIMS (positive mode) *m/z* calculated for C_27_H_23_O_7_^+^ [M + H]^+^ = 459.1438; observed [M + H]^+^ = 459.1435; Δ = -1.8594 ppm

Fasamycin C **13**: yield 51.68mg; deep yellow powder; UV (PDA) λ_max_: 245, 290, 355, 420 nm; IR (neat) *v*_max_ (cm^−1^) 3338, 2947, 2834, 1644, 1449, 1202, 1114, 1019, 617; ^1^H, ^13^C NMR data, see Table S3; Molecular formula: C_28_H_24_O_7_ ; HRESIMS (positive mode) *m/z* calculated for C_28_H_25_O_7_^+^ [M + H]^+^ = 473.1595; observed [M + H]^+^ = 473.1598; Δ = -1.0833ppm

### Minimum Inhibitory Concentration

Minimum inhibitory concentrations (MIC) of compounds **1-13** were determined against a range of Gram-positive bacteria, *S. aureus* (ATCC 25923), *E. faecalis* (ATCC 29212) and Gram-negative bacteria, *E. coli* (ATCC 25922), *P. aeruginosa* (ATCC 27853), fungal pathogen, *Candida albicans* ATCC 10231 as well as clinical strains of *E. faecium* K59-68 and *E. faecium* K60-39, and *S. haemolyticus* 8-7A. The hospital *E. faecium* isolates (K59-68 and K60-39) were isolated from the bloodstream of patients at the University Hospital of North Norway [27], and belonged to the complex clonal 17 sub-cluster which are highly resistant to ampicillin [27,28]. *Staphylococcus haemolyticus* 8-7A was obtained from the same hospital. All clinical strains were provided courtesy of Prof. Kristin Hegstad. The activity of **1-13** was determined using a sequential 2-fold serial dilution of the compounds (50-0.10 µg/mL) in DMSO following the standard protocols recommended by the Clinical and Laboratory Standard Institute [29] and as previously described [25,30,31]. The MIC was defined as the lowest concentration of the compound that inhibited ≥ 95% bacterial growth after overnight incubation. Ampicillin (Sigma) was used as the antibiotic standard.

## Author Contributions

Formal analysis and investigation, FM and YZ; data curation, FM; writing— original draft preparation, FM and HD; writing—review and editing, FM, HD and KK; supervision, HD; project administration, HD; funding acquisition, HD and KK. All authors have read and agreed to the published version of the manuscript.

## Acknowledgments

FM is grateful to Shan Wang, Ming Him Tong, and Lin Rui Wu for the protocols in gene deletion

