## Supplemental information for "Discovery of new antibacterial accramycins from a genetic variant of the soil bacterium, *Streptomyces* sp. MA37"

**List of Tables**

Table S1. Primers used in the Study

Table S2. Physico-chemical Properties of Accramycins A-K **1-11**

Table S3. ^1^H and ^13^C of Accramycins A-K **1-11**, naphthacemycin B1 **12** and fasamycin C **13**

(CD_3_OD, 298K, 600MHz)

**List of Figures**

Figure S1. HPLC traces of *Streptomyces* sp. MA37 mutant strain compared to Wild Type

Figure S2. Key HMBC correlations (→) of accramycins A-K **1-11**, naphthacemycin B1 **12**,

and fasamycin C **13** (CD_3_OD, 298K, 600MHz)

Figure S3. HRESIMS of naphthacemycin B1 **12**

Figure S4. ^1^H-NMR of naphthacemycin B1 **12** (CD_3_OD, 298K, 600MHz)

Figure S5. ^1^H-1H COSY of naphthacemycin B1 **12**(CD_3_OD, 298K, 600MHz)

Figure S6. HSQC of naphthacemycin B1 **12** (CD_3_OD, 298K, 600MHz)

Figure S7. HMBC of naphthacemycin B1 **12** (CD_3_OD, 298K, 600MHz)

Figure S8. NOESY of naphthacemycin B1 **12** (CD_3_OD, 298K, 600MHz)

Figure S9. HRESIMS of fasamycin C **13**

Figure S10. ^1^H-NMR of fasamycin C **13** (CD_3_OD, 298K, 600MHz)

Figure S11. ^1^H-^1^H COSY of fasamycin C **13** (CD_3_OD, 298K, 600MHz)

Figure S12. HSQC of fasamycin C **13** (CD_3_OD, 298K, 600MHz)

Figure S13. HMBC of fasamycin C **13** (CD_3_OD, 298K, 600MHz)

Figure S14. NOESY of fasamycin C **13** (CD_3_OD, 298K, 600MHz)

Figure S15. HRESIMS of Accramycin A **1**

Figure S16. ^1^H-NMR of Accramycin A **1** (CD_3_OD, 298K, 600MHz)

Figure S17. ^1^H-^1^H COSY of Accramycin A **1** (CD_3_OD, 298K, 600MHz)

Figure S18. HSQC of Accramycin A **1** (CD_3_OD, 298K, 600MHz)

Figure S19. HMBC of Accramycin A **1** (CD_3_OD, 298K, 600MHz)

Figure S20. NOESY of Accramycin A **1** (CD_3_OD, 298K, 600MHz)

Figure S21. HRESIMS of Accramycin B **2**

Figure S22. ^1^H-NMR of Accramycin B **2** (CD_3_OD, 298K, 600MHz)

Figure S23. ^1^H-^1^H COSY of Accramycin B **2** (CD_3_OD, 298K, 600MHz)

Figure S24. HSQC of Accramycin B **2** (CD_3_OD, 298K, 600MHz)

Figure S25. HMBC of Accramycin B **2** (CD_3_OD, 298K, 600MHz)

Figure S26. **A.** HRESIMS and **B**. Isotope Pattern of Accramycin C **3**

Figure S27. ^1^H-NMR of Accramycin C **3** (CD_3_OD, 298K, 600MHz)

Figure S28. ^1^H-^1^H COSY of Accramycin C **3** (CD_3_OD, 298K, 600MHz)

Figure S29. HSQC of Accramycin C **3** (CD_3_OD, 298K, 600MHz)

Figure S30. NOESY of Accramycin C **3** (CD_3_OD, 298K, 600MHz)

Figure S31. **A**. HRESIMS and **B**. Isotope Pattern of Accramycin D **4**

Figure S32. ^1^H-NMR of Accramycin D **4** (CD_3_OD, 298K, 600MHz)

Figure S33. ^1^H-^1^H COSY of Accramycin D **4** (CD_3_OD, 298K, 600MHz)

Figure S34. HSQC of Accramycin D **4** (CD_3_OD, 298K, 600MHz)

Figure S35. HMBC of Accramycin D **4** (CD_3_OD, 298K, 600MHz)

Figure S36. **A.** HRESIMS and **B.** Isotope Pattern of Accramycin E **5**

Figure S37. ^1^H-NMR of Accramycin E **5** (CD_3_OD, 298K, 600MHz)

Figure S38. ^1^H-^1^H COSY of Accramycin E **5** (CD_3_OD, 298K, 600MHz)

Figure S39. HSQC of Accramycin E **5** (CD_3_OD, 298K, 600MHz)

Figure S40. HMBC of Accramycin E **5** (CD_3_OD, 298K, 600MHz)

Figure S41. **A.** HRESIMS and **B.** Isotope Pattern of Accramycin F **6**

Figure S42. ^1^H-NMR of Accramycin F **6** (CD_3_OD, 298K, 600MHz)

Figure S43. ^1^H-^1^H COSY of Accramycin F **6** (CD_3_OD, 298K, 600MHz)

Figure S44. HSQC of Accramycin F **6** (CD_3_OD, 298K, 600MHz)

Figure S45. HMBC of Accramycin F **6** (CD_3_OD, 298K, 600MHz)

Figure S46. NOESY of Accramycin F **6** (CD_3_OD, 298K, 600MHz)

Figure S47. **A.** HRESIMS and **B.** Isotope Pattern of Accramycin G **7**

Figure S48. ^1^H-NMR of Accramycin G **7** (CD_3_OD, 298K, 600MHz)

Figure S49. ^1^H-^1^H COSY of Accramycin G **7** (CD_3_OD, 298K, 600MHz)

Figure S50. HSQC of Accramycin G **7** (CD_3_OD, 298K, 600MHz)

Figure S51. HMBC of Accramycin G **7** (CD_3_OD, 298K, 600MHz)

Figure S52. NOESY of Accramycin G **7** (CD_3_OD, 298K, 600MHz)

Figure S53. **A.** HRESIMS and **B.** Isotope Pattern of Accramycin H **8**

Figure S54. ^1^H-NMR of Accramycin H **8** (CD_3_OD, 298K, 600MHz)

Figure S55. ^1^H-^1^H COSY of Accramycin H **8** (CD_3_OD, 298K, 600MHz)

Figure S56. HSQC of Accramycin H **8** (CD_3_OD, 298K, 600MHz)

Figure S57. HMBC of Accramycin H **8** (CD_3_OD, 298K, 600MHz)

Figure S58. **A.** HRESIMS and **B.** Isotope Pattern of Accramycin I **9**

Figure S59. ^1^H-NMR of Accramycin I **9** (CD_3_OD, 298K, 600MHz)

Figure S60. ^1^H-^1^H COSY of Accramycin I **9** (CD_3_OD, 298K, 600MHz)

Figure S61. HSQC of Accramycin I **9** (CD_3_OD, 298K, 600MHz)

Figure S62. HMBC of Accramycin I **9** (CD_3_OD, 298K, 600MHz)

Figure S63. NOESY of Accramycin I **9** (CD_3_OD, 298K, 600MHz)

Figure S64. LCMS isotope pattern of Accramycin J **10**

Figure S65. ^1^H-NMR of Accramycin J **10** (CD_3_OD, 298K, 600MHz)

Figure S66. ^1^H-^1^H COSY of Accramycin J **10** (CD_3_OD, 298K, 600MHz)

Figure S67. HSQC of Accramycin J **10** (CD_3_OD, 298K, 600MHz)

Figure S68. HMBC of Accramycin J **10** (CD_3_OD, 298K, 600MHz)

Figure S69. NOESY of Accramycin J **10** (CD_3_OD, 298K, 600MHz)

Figure S70. LCMS Isotope Pattern of Accramycin K **11**

Figure S71. ^1^H-NMR of Accramycin K **11** (CD_3_OD, 298K, 600MHz)

Figure S72. ^1^H-^1^H COSY of Accramycin K **11** (CD_3_OD, 298K, 600MHz)

Figure S73. HSQC of Accramycin K **11** (CD_3_OD, 298K, 600MHz)

Figure S74. HMBC of Accramycin K **11** (CD_3_OD, 298K, 600MHz)

Figure S75. NOESY of Accramycin K **11** (CD_3_OD, 298K, 600MHz)

Table S1. Primers used in the Study

| **Primer ID** | **Primer Sequence** | **Purpose** |
| --- | --- | --- |
| MarR-FRA | CATGACCTCTAGACTCAAGAAGGCCTCCGC GAACTGA | Right Arm Forward: Construction of knockout vector |
| MarR-RRA | ACATGATTACGAATTCGGATGCGCTGGGTG CAGGACTT | Right Arm Reverse: Construction of knockout vector |
| MarR-FLA | GGCCAGTGCCAAGCTTTCCCGCCATGCACA CCACACTGAT | Left Arm Forward: Construction of knockout vector |
| MarR-RLA | TCTTGAGTCTAGAGGTCATGGTCGGTCACC TCTGCCCT | Left Arm Reverse: Construction of knockout vector |
| LuxR-FRA | AATTCGTAATCATGTCATAGCTGTTTCCTGTG | Right Arm Forward: Construction of knockout vector |
| LuxR-RRA | ACATGATTACGAATTTCATCCGCGACCGG  ATCGA | Right Arm Reverse: Construction of knockout vector |
| LuxR-FLA | GGCCAGTGCCAAGCTGTGAGCGGGGGC  TGCGC | Left Arm Forward: Construction of knockout vector |
| LuxR-RLA | TGGCACTGGCCGTCGTTTTACAACGTCGTGAC TGGG | Left Arm Reverse: Construction of knockout vector |
| LysR-FRA | AATTCGTAATCATGTCATAGCTGTTTCCTGTG | Right Arm Forward: Construction of knockout vector |
| LysR-RRA | ACATGATTACGAATTTCAGGTGTCGGC  GCCG | Right Arm Reverse: Construction of knockout vector |
| LysR-FLA | GGCCAGTGCCAAGCTATGGAGCTCCGGCA  GCTGCA | Left Arm Forward: Construction of knockout vector |
| LysR-RLA | TGGCACTGGCCGTCGTTTTACAACGTC GTGACTG | Left Arm Reverse: Construction of knockout vector |
| MerR-FRA | AATTCGTAATCATGTCATAGCTGTTTCCTGTG | Right Arm Forward: Construction of knockout vector |
| MerR-RRA | ACATGATTACGAATTTCACCCGGTGGTGATCG  GCTCCTG | Right Arm Reverse: Construction of knockout vector |
| MerR-FLA | CGACGGCCAGTGCCAATGTTCAGTATCGGAG  ACTTCGC | Left Arm Forward: Construction of knockout vector |
| MerR-RLA | TGGCACTGGCCGTCGTTTTACAACGTCGT  GACTGGG | Left Arm Reverse: Construction of knockout vector |
| Mu-F | CGCACCCTTCTGTCGGACCACCACTGA | Mutant verification |
| Mu-R | GTTCTGCTTGGCGACGCTGACCGAGTA | Mutant verification |

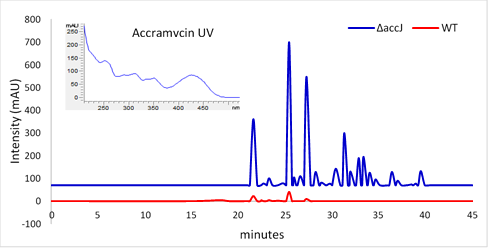

Figure S1. HPLC traces monitored at λ450nm of Streptomyces sp. MA37 mutant strain (blue) compared to Wild Type (red) (5µL injection, 5mg/mL) with the characteristic accramycin UV spectrum

Table S2. Physico-chemical Properties of Accramycins A-K **1-11**, naphthacemycin B1 **12**, and fasamycin C **13** from *Streptomyces* sp. MA37

|  | Accramycin A **1** | Accramycin B **2** | Accramycin C **3** | Accramycin D **4** |
| --- | --- | --- | --- | --- |
| Appearance | deep yellow powder | deep yellow powder | deep yellow powder | deep yellow powder |
| Molecular formula | C_29_H_26_O_7_ | C_30_H_28_O_7_ | C_28_H_23_ClO_7_ | C_30_H_27_ClO_7_ |
| HR ESIMS (*obs*) | 487.1748 [M+H]^+^ | 501.1898 [M+H]^+^ | 507.1207 [M+H]^+^ | 535.1511 [M+H]^+^ |
| *m/z* (*calc*) | 487.1751 (for C_29_H_27_O_7_^+^) | 501.1912 (for C_30_H_29_O_7_^+^) | 507.1205 (for C_28_H_24_ClO_7_^+^) | 535.1518 (for C_30_H_28_ClO_7_^+^) |
| ∆ ppm | -1.78 | -1.98 | 0.37 | -1.39 |
| IR v_max_ (cm^-1^) | 3350, 2946, 2834, 1681, 1607, 1284, 1202, 1026, 584 | 3350, 2946, 2834, 1681, 1607, 1284, 1202, 1026, 584 | 3362, 2922, 2848, 1679, 1612, 1443, 1203, 1149, 726 | 3337, 2947, 2834, 1681, 1450, 1025, 634 |
| UV (PDA) λ_max_ | 225, 245, 290, 355, 420 | 250, 290, 315, 335, 350, 425 | 250, 280, 300, 355, 430 | 250, 290, 315, 335, 350, 425 |

|  | Accramycin E **5** | Accramycin F **6** | Accramycin G **7** | Accramycin H **8** |
| --- | --- | --- | --- | --- |
| Appearance | deep yellow powder | deep yellow powder | deep yellow powder | deep yellow powder |
| Molecular formula | C_29_H_24_Cl_2_O_7_ | C_29_H_24_Cl_2_O_7_ | C_30_H_26_Cl_2_O_7_ | C_28_H_21_Cl_3_O_7_ |
| HR ESIMS (*obs*) | 555.0978 [M+H]^+^ | 555.0975 [M+H]^+^ | 569.1127 [M+H]^+^ | 575.0420 [M+H]^+^ |
| *m/z* (*calc*) | 555.0972 (for C_29_H_25_Cl_2_O_7_^+^) | 555.0972 (for C_29_H_25_Cl_2_O_7_^+^) | 569.1128 (for C_30_H_27_Cl_2_O_7_^+^) | 575.0426 (for C_28_H_22_Cl_3_O_7_^+^) |
| ∆ ppm | 1.18 | 0.63 | -0.29 | -0.99 |
| IR v_max_ (cm^-1^) | 3337, 2947, 2834, 1681, 1450, 1025, 634 | 3325, 2943, 2833, 1678, 1449, 1119, 1023, 635 | 3399, 2917, 2851, 1688, 1606, 1423, 1322, 1205 | 3386, 1682, 1439, 1327, 1197, 1136, 844, 802, 725 |
| UV (PDA) λ_max_ | 250, 290, 315, 335, 350, 425 | 250, 290, 315, 335, 350, 425 | 250, 295, 320, 340, 350, 425 | 250, 290, 310, 355, 425 |

Table S2. Physico-chemical Properties of Accramycins A-K **1-11**, naphthacemycin B1 **12**, and fasamycin C **13** from *Streptomyces* sp. MA37

|  | Accramycin I **9** | Accramycin J **10** | Accramycin K **11** |
| --- | --- | --- | --- |
| Appearance | deep yellow powder | deep yellow powder | deep yellow powder |
| Molecular formula | C_29_H_23_Cl_3_O_7_ | C_28_H_20_Cl_4_O_7_ | C_29_H_22_Cl_4_O_7_ |
| HR ESIMS (*obs*) | 589.0574 [M+H]^+^ | 609.0040 [M+H]^+^ | 623.0184 [M+H]^+^ |
| *m/z* (*calc*) | 589.0582 (for C_29_H_24_Cl_3_O_7_^+^) | 609.0036 (for C_28_H_21_Cl_4_O_7_^+^) | 623.0192 (for C_29_H_23_Cl_4_O_7_^+^) |
| ∆ ppm | -1.32 | 0.72 | 0.41 |
| IR v_max_ (cm^-1^) | 3399, 2921, 2849, 1680, 1442, 1196, 1139 | 3427, 1688, 1601, 1439, 1328, 1204, 1139 | 3427, 1688, 1601, 1439, 1328, 1204, 1139 |
| UV (PDA) λ_max_ | 250, 290, 315, 340, 350, 425 | 250, 290, 315, 340, 350, 425 | 250, 295, 320, 340, 350, 425 |

|  | Naphthacemycin B1 **12** | Fasamycin C **13** |
| --- | --- | --- |
| Appearance | reddish powder | deep yellow powder |
| Molecular formula | C_27_H_22_O_7_ | C_28_H_24_O_7_ |
| HR ESIMS (*obs*) | 459.1435 [M+H]^+^ | 473.1598 [M+H]^+^ |
| *m/z* (*calc*) | 459.1438 (for C_27_H_23_O_7_^+^) | 473.1595 (for C_28_H_25_O_7_^+^) |
| ∆ ppm | -1.86 | -1.08 |
| IR v_max_ (cm^-1^) | 3337, 2946, 1678, 1448, 1204, 1021, 644 | 3338, 2947, 2834, 1644, 1449, 1202, 1114, 1019, 617 |
| UV (PDA) λ_max_ | 245, 290, 355, 420 | 245, 290, 355, 420 |

Table S3. ^1^H and ^13^C of Accramycins A-K **1-11**, naphthacemycin B1 **12** and fasamycin C **13** (CD_3_OD, 298K, 600MHz)

| **no.** | Accramycin A **1** | | Accramycin B **2** | | Accramycin C **3** | | Accramycin D **4** | | Accramycin E **5** | |
| --- | --- | --- | --- | --- | --- | --- | --- | --- | --- | --- |
|  | ^13^C | ^1^H, mult. (J,Hz) | ^13^C | ^1^H, mult. (J,Hz) | ^13^C | ^1^H, mult. (J,Hz) | ^13^C | ^1^H, mult. (J,Hz) | ^13^C | ^1^H, mult. (J,Hz) |
| 1 | 105.9, CH | 6.67, d (2.4) | 106.1, CH | 6.66, d (1.4) | 105.4, CH | 6.84, s | 101.0, CH | 6.97, s | 105.7, CH | 6.87, s |
| 2 | 165.7, C | - | 166.5, C | - |  | - | 161.2, C | - | 160.7, C | - |
| 3 | 100.9, CH | 6.22, d (2.4) | 98.7, CH | 6.42, d (2.0) |  | - | 106.3, C | - | 106.8, C | - |
| 4 | 165.7, C | - | 165.8, C | - |  | - | 161.3, C | - | 160.7, C | - |
| 5 | 107.5, C | - | 108.0, C | - |  | - | 108.4, C | - | 108.4, C | - |
| 6 | 190.4, C | - | 190.8, C | - |  | - | 190.4, C | - | 190.4, C | - |
| 7 | 107.4, C | - | 106.2, C | - |  | - | 106.3, C | - | 106.6, C | - |
| 8 | 165.3, C | - | 165.0, C | - |  | - | 165.7, C | - | 165.7, C | - |
| 9 | 117.4, C | - | 117.6, C | - |  | - | 116.9, C | - | 117.4, C | - |
| 10 | 141.0, C | - | 141.0, C | - |  | - | 141.9, C | - | 137.3, C | - |
| 11 | 121.7, CH | 6.75, d (2.4) | 121.4, CH | 6.77, d (2.1) | 121.1, CH | 6.73, d (2.1) | 121.4, CH | 6.77, d (2.1) | 115.9, CH | 7.05, d (2.1) |
| 12 | 161.1, C | - | 160.9, C |  |  | - | 161.2, C | - | 155.8, C | - |
| 13 | 105.8, CH | 7.21, d (2.4) | 105.8, CH | 7.25, d (2.0) | 109.0, CH | 7.08, d (2.1) | 105.9, CH | 7.25, d (2.1) | 115.2, C | - |
| 14 | 141.8, C | - | 141.3, C |  |  | - | 141.9, C | - | 138.4, C | - |
| 15 | 115.8, CH | 7.51, s | 115.5, CH | 7.56, s | 115.0, CH | 7.39, s | 115.8, CH | 7.57, s | 111.2, CH | 7.95, s |
| 16 | 145.3, C | - | 145.3, C | - | 144.8, C | - | 145.3, C | - | 146.4, C | - |
| 17 | 39.3, C | - | 38.6, C | - | 38.7, C | - | 38.4, C | - | 38.4, C | - |
| 18 | 154.6, C | - | 154.0, C | - | 151.9, C | - | 152.4, C | - | 151.7, C | - |
| 19 | 34.4, CH_3_ | 1.70, s | 33.3, CH_3_ | 1.77, s | 33.1, CH_3_ | 1.72, s | 33.4, CH_3_ | 1.80, s | 33.4, CH_3_ | 1.77, s |
| 20 | 34.7, CH_3_ | 1.69, s | 33.4, CH_3_ | 1.75, s | 33.1, CH_3_ | 1.71, s | 33.4, CH_3_ | 1.79, s | 33.4, CH_3_ | 1.75, s |
| 21 | 123.9, C | - | 124.4 C | - | 124.5, C | - | 123.9, C | - | 123.9, C | - |
| 22 | 154.2, C | - | 154.5, C | - |  | - | 154.3, C | - | 154.3, C | - |
| 23 | 98.3, CH | 6.33, d (2.4) | 98.3, CH | 6.35, s | 98.3, CH | 6.33, d (2.1) | 98.4, CH | 6.34, d (2.1) | 98.4, CH | 6.35, d (2.1) |
| 24 | 159.1, C | - | 159.2,C | - | 158.9, C | - | 159.5, C | - | 159.3, C | - |
| 25 | 105.9, CH | 6.37, d (2.4) | 106.0, CH | 6.40, s | 106.5, CH | 6.38, d (2.1) | 105.9, CH | 6.38, d (2.1) | 105.9, CH | 6.40, d (2.1) |
| 26 | 137.0, C | - | 136.9, C |  | 136.8, C | - | 136.8, C | - | 136.8, C | - |
| 27 | 20.6, CH_3_ | 1.91, s | 19.5, CH_3_ | 1.93, s | 19.5, CH_3_ | 1.92, s | 19.4, CH_3_ | 1.91, s | 19.4, CH_3_ | 1.93, s |
| 28 | 55.3, CH_3_ | 3.80, s | 54.3, CH_3_ | 3.83, s | 54.3, CH_3_ | 3.80, s | 54.2, CH_3_ | 3.81, s | 54.2, CH_3_ | 3.81, s |
| 29 | 55.6, CH_3_ | 3.95, s | 54.7, CH_3_ | 3.99, s | - | - | 54.7, CH_3_ | 3.98, s | 55.7, CH_3_ | 4.02, s |
| 30 | - | - | 54.9, CH_3_ | 3.91, s | - | - | 55.8, CH_3_ | 4.05, s | - | - |

Table S3. ^1^H and ^13^C of Accramycins A-K **1-11**, naphthacemycin B1 **12** and fasamycin C **13** (CD_3_OD, 298K, 600MHz)

| **no.** | Accramycin F **6** | | Accramycin G **7** | | Accramycin H **8** | | Accramycin I **9** | | Accramycin J **10** | |
| --- | --- | --- | --- | --- | --- | --- | --- | --- | --- | --- |
|  | ^13^C | ^1^H, mult. (J,Hz) | ^13^C | ^1^H, mult. (J,Hz) | ^13^C | ^1^H, mult. (J,Hz) | ^13^C | ^1^H, mult. (J,Hz) | ^13^C | ^1^H, mult. (J,Hz) |
| 1 | 101.5, CH | 7.00, s | 101.3, CH | 7.00, s | 101.1, CH | 6.97, s | 101.5, CH | 7.00, s | 101.5, CH | 7.00, s |
| 2 | 161.5, C | - | 161.9, C | - | 161.3, C | - | 161.6, C | - | 161.9, C | - |
| 3 | 107.6, C | - | 108.1, C | - | 108.8, C | - | 108.1, C | - | 108.0, C | - |
| 4 | 161.7, C | - | 161.7, C | - | 160.8, C | - | 161.8, C | - | 162.0, C | - |
| 5 | 108.1, C | - | 108.1, C | - | 108.8, C | - | 108.1, C | - | 108.0, C | - |
| 6 | 190.6, C | - | 190.6, C | - | 190.0, C | - | 190.5, C | - | 190.7, C | - |
| 7 | 106.8, C | - | 106.7, C | - | 106.6, C | - | 107.0, C | - | 107.2, C | - |
| 8 | 165.7, C | - | 166.0, C | - | 165.5, C | - | 164.3, C | - | 165.0, C | - |
| 9 | 117.7, C | - | 118.0, C | - | 116.8, C | - | 118.0, C | - | 117.7, C | - |
| 10 | 137.3, C | - | 138.5, C | - | 141.3, C | - | 138.4, C | - | 137.1, C | - |
| 11 | 120.9, CH | 6.87, d (2.1) | 116.2, CH | 7.07, s | 121.3, CH | 6.74, s | 116.3, CH | 7.06, s | 120.9, CH | 6.87, s |
| 12 | 154.9, C | - | 156.6, C | - | 163.5, C | - | 155.9, C | - | 155.0, C | - |
| 13 | 112.9, C | - | 115.0, C | - | 109.4, CH | 7.14, d (2.1) | 115.5, C | - | 113.8, C | - |
| 14 | 138.4, C | - | 146.5, C | - | 142.5, C | - | 138.4, C | - | 138.4, C | - |
| 15 | 111.2, CH | 7.93, s | 111.3, CH | 7.98, s | 115.2, CH | 7.45, s | 111.3, CH | 7.99, s | 111.7, CH | 7.95, s |
| 16 | 146.7, C | - | 146.5, C | - | 148.5, C | - | 146.9, C | - | 146.8, C | - |
| 17 | 39.4, C | - | 39.1, C | - | 39.4, C | - | 39.5, C | - | 39.6, C | - |
| 18 | 152.5, C | - | 152.5, C | - | 153.2, C | - | 152.5, C | - | 152.5, C | - |
| 19 | 33.4, CH_3_ | 1.83, s | 33.4, CH_3_ | 1.83, s | 33.2, CH_3_ | 1.79, s | 33.5, CH_3_ | 1.83, s | 33.5, CH_3_ | 1.83, s |
| 20 | 33.4, CH_3_ | 1.82, s | 33.4, CH_3_ | 1.82, s | 33.2, CH_3_ | 1.78, s | 33.5, CH_3_ | 1.82, s | 33.5, CH_3_ | 1.82, s |
| 21 | 124.2, C | - | 123.7, C | - | 125.5, C | - | 124.5, C | - | 124.9, C | - |
| 22 | 159.5, C | - | 159.4, C | - | 169.0, C | - | 152.5, C | - | 152.5, C | - |
| 23 | 98.1, CH | 6.32, d (2.3) | 98.3, CH | 6.34, d (2.3) | 107.5, C | - | 100.7, CH | 6.44, s | 107.5, CH | - |
| 24 | 159.3, C | - | 159.4, C | - | 169.0, C | - | 152.5, C | - | 148.4, C | - |
| 25 | 106.3, CH | 6.38, d (2.3) | 105.9, CH | 6.40, d (2.3) | 113.2, C | - | 112.1, C | - | 113.2, C | - |
| 26 | 137.2, C | - | 137.0, C | - | 133.1, C | - | 134.7, C | - | 132.5, C | - |
| 27 | 19.4, CH_3_ | 1.93, s | 19.4, CH_3_ | 1.93, s | 16.9, CH_3_ | 1.93, s | 17.1, CH_3_ | 1.98, s | 16.9, CH_3_ | 1.98, s |
| 28 | 54.4, CH_3_ | 3.80, s | 54.2, CH_3_ | 3.81, s | - | - | - | - | - | - |
| 29 | - | - | 55.7, CH_3_ | 4.02, s | - | - | 56.0, CH_3_ | 4.02, s | - | - |
| 30 | 55.8, CH_3_ | 4.06, s | 55.7, CH_3_ | 4.06, s | 55.7, CH_3_ | 4.05, s | 55.7, CH_3_ | 4.06, s | 56.0, CH_3_ | 4.06, s |

Table S3. ^1^H and ^13^C of Accramycins A-K **1-11**, naphthacemycin B1 **12** and fasamycin C **13** (CD_3_OD, 298K, 600MHz)

| **no.** | Accramycin K **11** | | Naphthacemycin B1 **1**2 | | Fasamycin C **13** | |
| --- | --- | --- | --- | --- | --- | --- |
|  | ^13^C | ^1^H, mult. (J,Hz) | ^13^C | ^1^H, mult. (J,Hz) | ^13^C | ^1^H, mult. (J,Hz) |
| 1 | 101.4, CH | 7.00, s | 106.6, CH | 6.66, d (2.1) | 106.1, CH | 6.66, d (2.3) |
| 2 | 161.5, C | - | 166.1, C | - | 165.9, C | - |
| 3 | 108.3, C | - | 101.2, CH | 6.21, d (2.1) | 101.2, CH | 6.21, d (2.0) |
| 4 | 161.8, C | - | 165.5, C | - | 165.5, C | - |
| 5 | 108.3, C | - | 108.8, C | - | 108.8, C | - |
| 6 | 190.6, C | - | 190.7, C | - | 190.4, C | - |
| 7 | 107.4, C | - | 107.4, C | - | 107.4, C | - |
| 8 | 165.6, C | - | 165.2, C | - | 165.2, C | - |
| 9 | 118.0, C | - | 116.7, C | - | 116.7, C | - |
| 10 | 137.4, C | - | 141.0, C | - | 141.0, C | - |
| 11 | 116.3, CH | 7.09, s | 121.7, CH | 6.72, d (2.2) | 121.7, CH | 6.72, d (2.1) |
| 12 | 156.2, C | - | 158.8, C | - | 158.8, C | - |
| 13 | 116.1, C | - | 109.3, CH | 7.06, d (2.5) | 109.3, CH | 7.06, d (2.0) |
| 14 | 141.4, C | - | 141.6, C | - | 141.6, C | - |
| 15 | 111.3, CH | 8.00, s | 115.3, CH | 7.36, s | 115.3, CH | 7.37, s |
| 16 | 148.6, C | - | 141.5 C | - | 141.5 C | - |
| 17 | 39.3, C | - | 38.7, C | - | 38.7, C | - |
| 18 | 152.4, C | - | 156.3, C | - | 156.3, C | - |
| 19 | 33.5, CH_3_ | 1.83, s | 34.0, CH_3_ | 1.71, s | 33.8, CH_3_ | 1.72, s |
| 20 | 33.5, CH_3_ | 1.82, s | 34.0, CH_3_ | 1.70, s | 33.8, CH_3_ | 1.70, s |
| 21 | 124.9, C | - | 124.4 C | - | 124.4 C | - |
| 22 | 152.4, C | - | 155.1, C | - | 155.1, C | - |
| 23 | 107.6, CH | - | 100.1, CH | 6.24, d (2.1) | 100.1, CH | 6.32, s |
| 24 | 148.6, C | - | 156.4, C | - | 159.3, C | - |
| 25 | 113.1, C | - | 107.9, CH | 6.27 ,d (2.3) | 107.9, CH | 6.37 , s |
| 26 | 132.4, C | - | 136.8, C | - | 136.8, C | - |
| 27 | 17.1, CH_3_ | 1.99, s | 19.8, CH_3_ | 1.87, s | 19.8, CH_3_ | 1.92, s |
| 28 | - | - | - | - | 54.0, CH_3_ | 3.80, s |
| 29 | 55.9, CH_3_ | 4.02, s | - | - | - | - |
| 30 | 55.7, CH_3_ | 4.06, s | - | - | - | - |

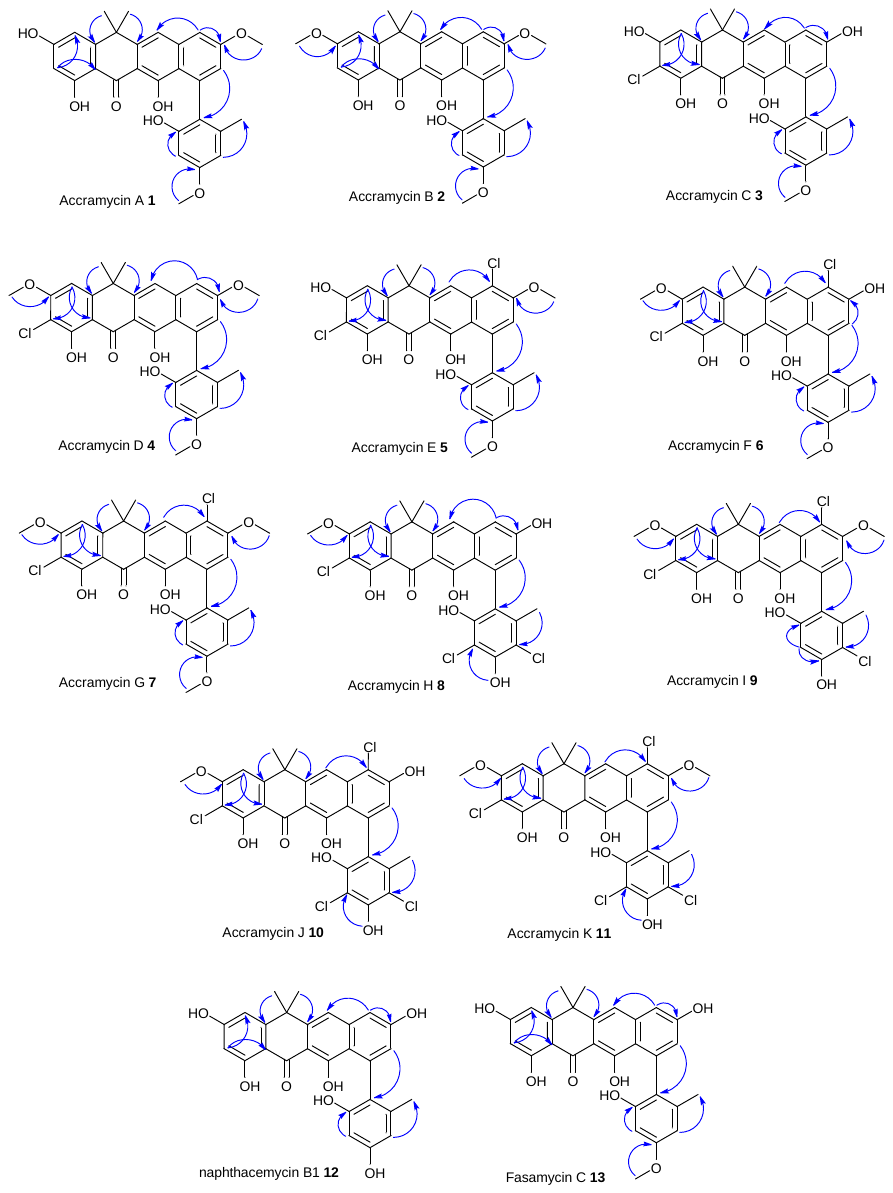

Figure S2. Key HMBC correlations (→) of accramycins A-K **1-11**, naphthacemycin B1 **12**, and fasamycin C **13** (CD_3_OD, 298K, 600MHz)

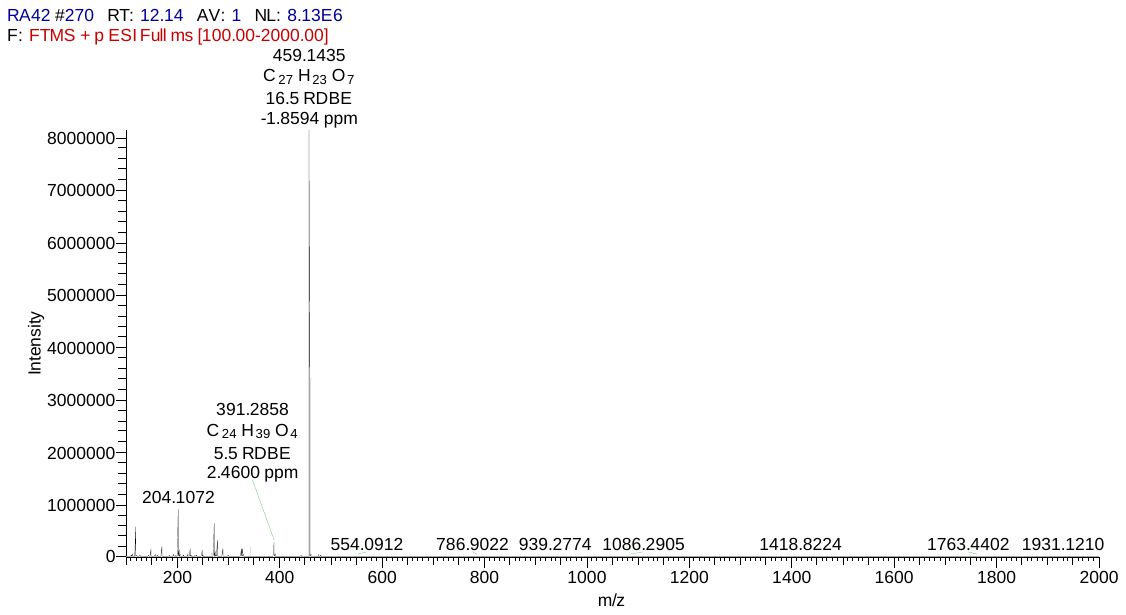

[M+H]^+^

Figure S3. HRESIMS of naphthacemycin B1 **12**

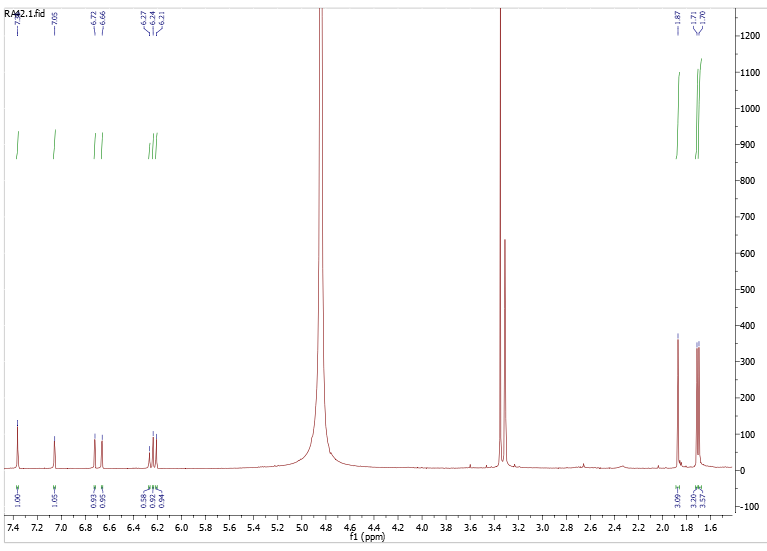

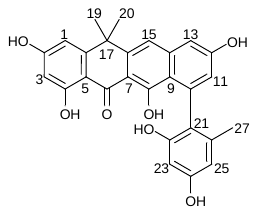

15

19, 20

27

13

1

11

25

23

3

Figure S4. ^1^H-NMR of naphthacemycin B1 **12** (CD_3_OD, 298K, 600MHz)

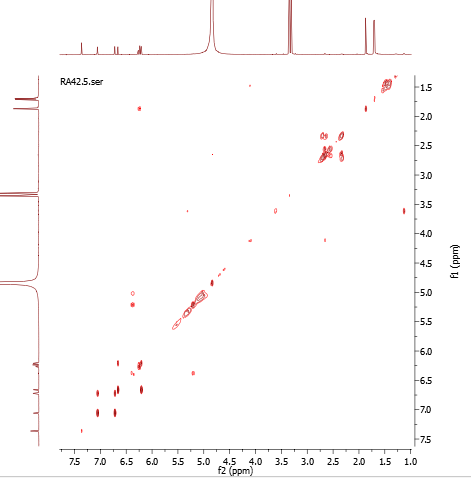

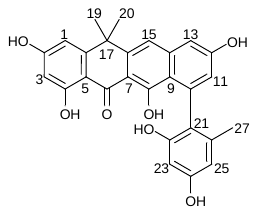

15

13

11

1

27

19, 20

25, 23, 3

19, 20

27

15

13

11

1

25, 23, 3

Figure S5. ^1^H-^1^H COSY of naphthacemycin B1 **12** (CD_3_OD, 298K, 600MHz)

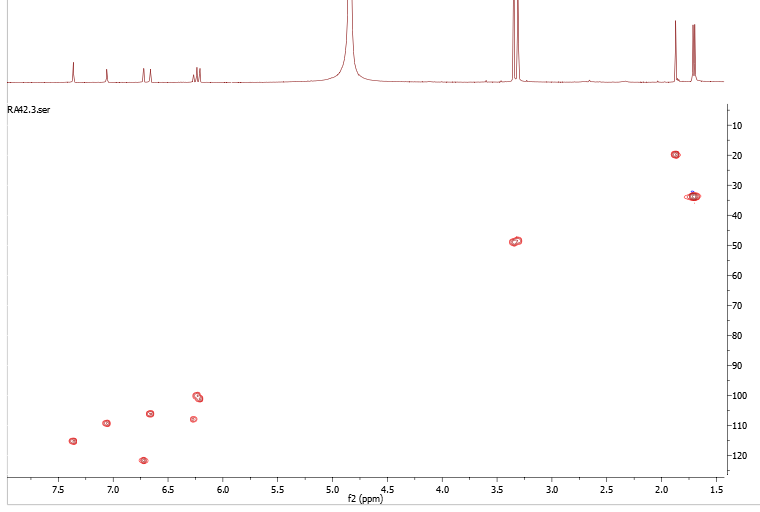

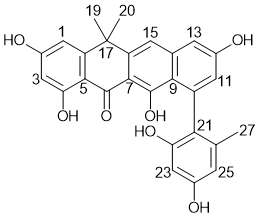

25, 23, 3

15

13

11

1

27

19, 20

Figure S6. HSQC of naphthacemycin B1 **12** (CD_3_OD, 298K, 600MHz)

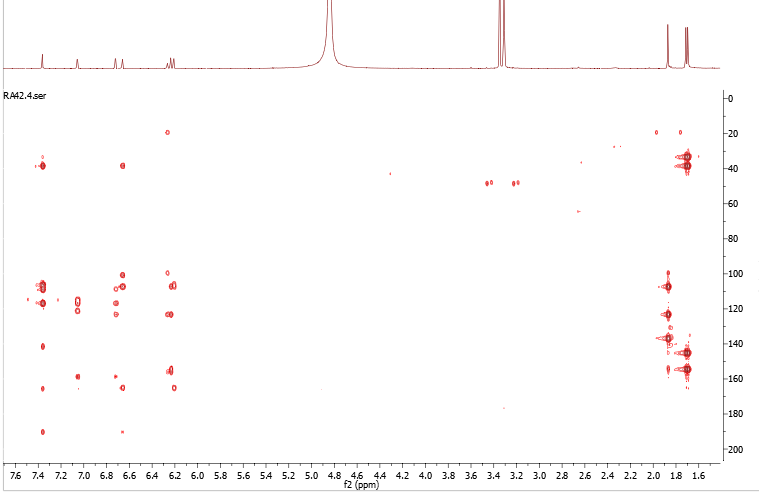

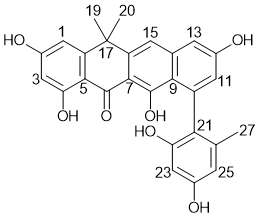

25, 23, 3

15

13

11

1

27

19, 20

Figure S7. HMBC of naphthacemycin B1 **12** (CD_3_OD, 298K, 600MHz)

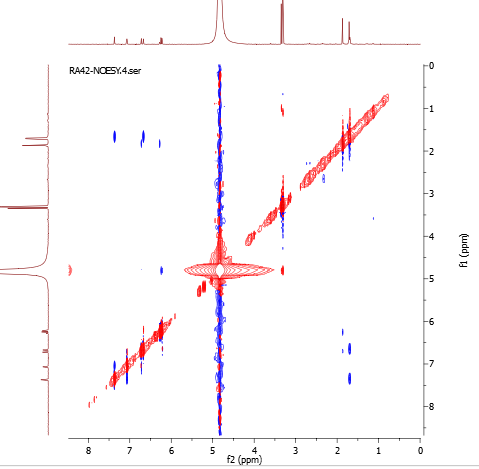

19, 20

27

19, 20

27

25, 23, 3

25, 23, 3

15

15

13

13

11, 1

11

1

Figure S8. NOESY of naphthacemycin B1 **12** (CD_3_OD, 298K, 600MHz)

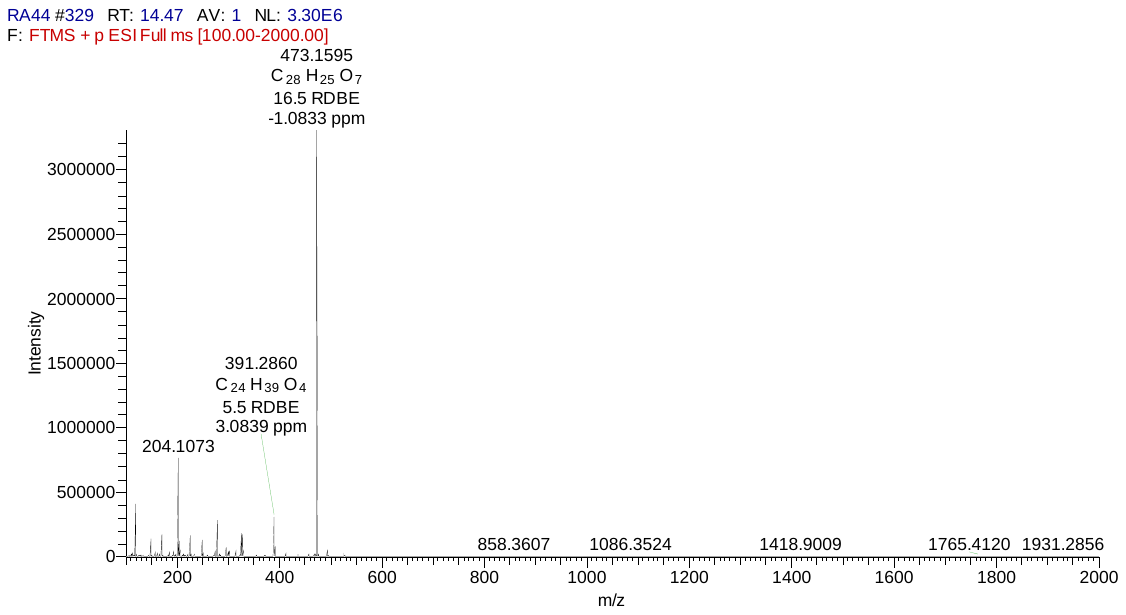

[M+H]^+^

Figure S9. HRESIMS of fasamycin C **13**

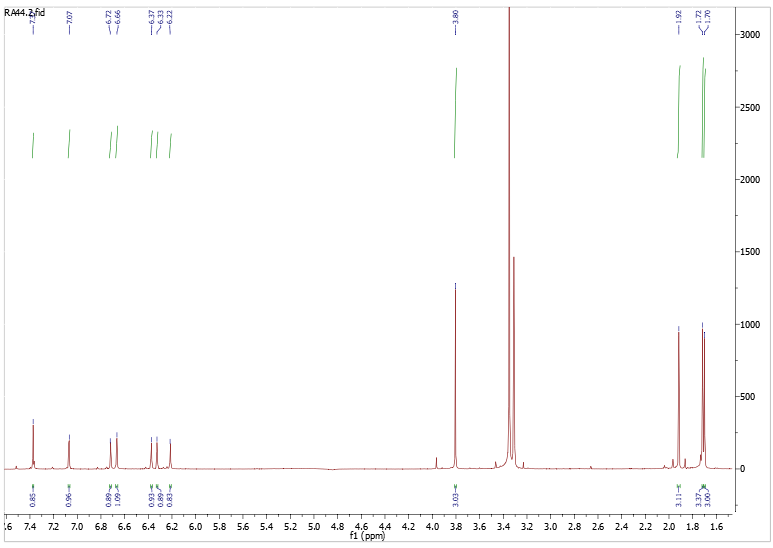

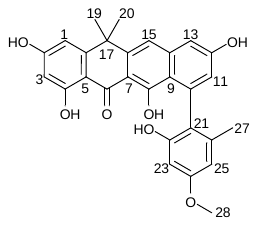

19, 20

27

15

13

11

1

25, 23, 3

28

Figure S10. ^1^H-NMR of fasamycin C **13** (CD_3_OD, 298K, 600MHz)

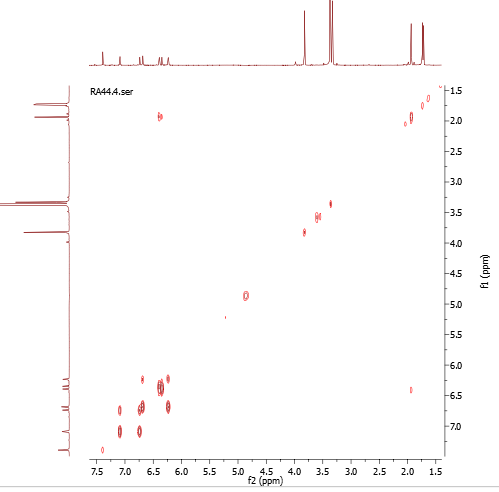

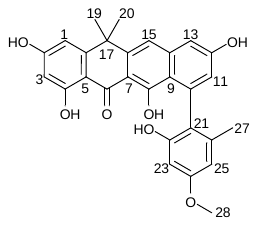

19, 20

19, 20

27

27

25, 23, 3

25, 23, 3

15

15

13

11

1

13

11

1

28

28

Figure S11. ^1^H-^1^H COSY of fasamycin C **13** (CD_3_OD, 298K, 600MHz)

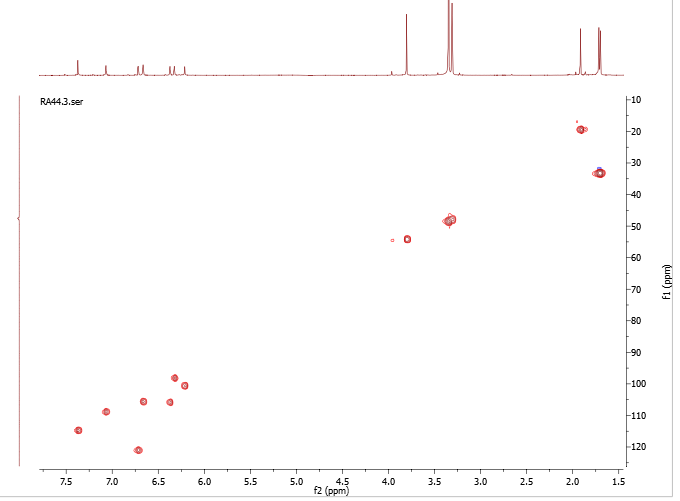

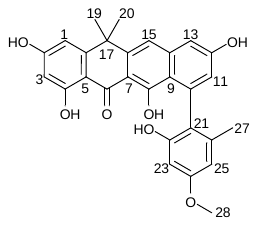

15

13

11

1

25

23

3

28

19, 20

27

Figure S12. HSQC of fasamycin C **13** (CD_3_OD, 298K, 600MHz)

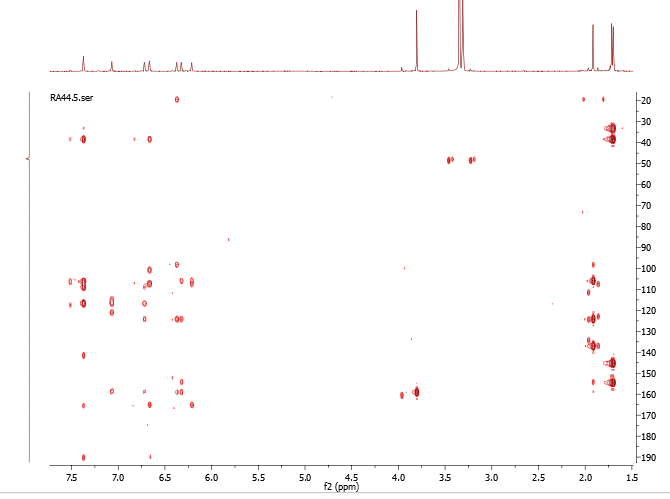

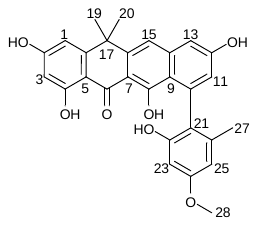

19, 20

27

28

15

13

11

1

25, 23, 3

Figure S13. HMBC of fasamycin C **13** (CD_3_OD, 298K, 600MHz)

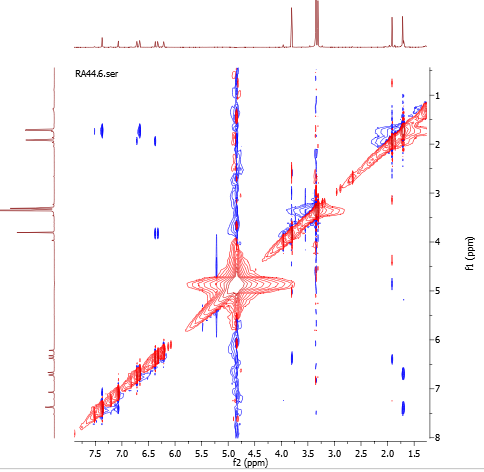

19, 20

19, 20

27

27

25, 23, 3

25, 23, 3

28

28

15

13

11

1

11

1

13

15

Figure S14. NOESY of fasamycin C **13** (CD_3_OD, 298K, 600MHz)

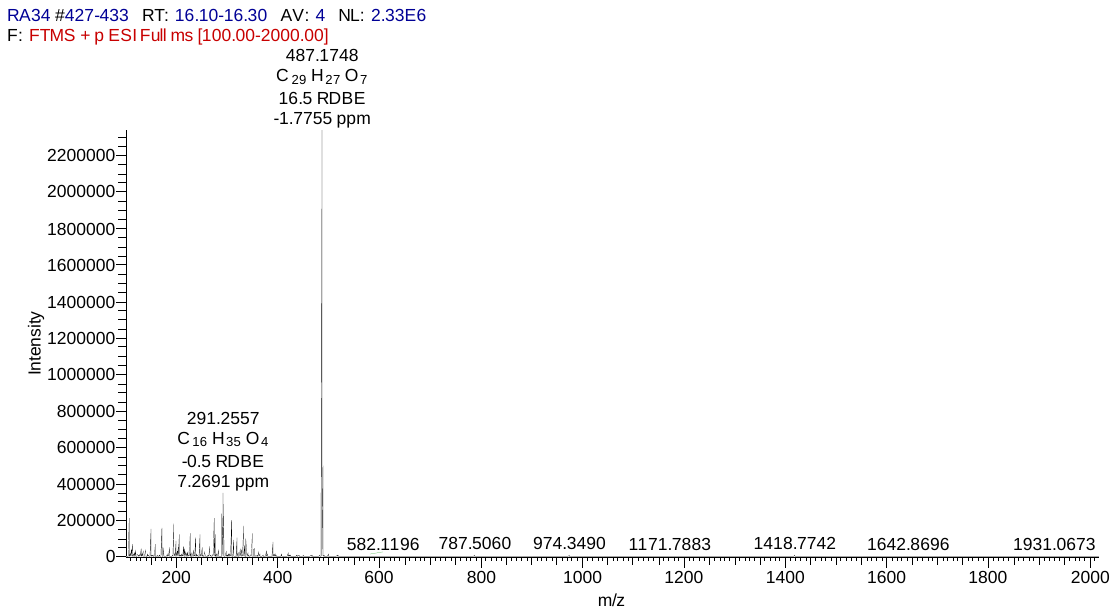

[M+H]^+^

Figure S15. HRESIMS of Accramycin A **1**

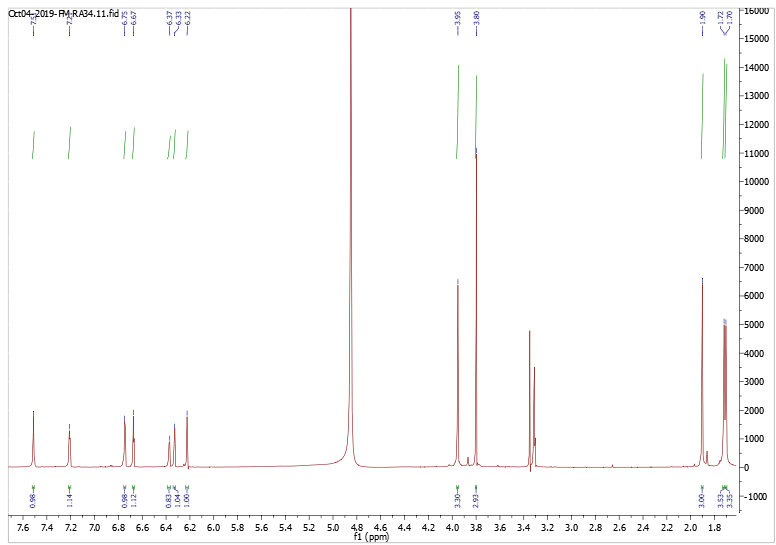

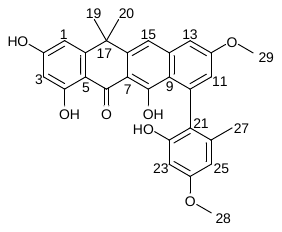

19, 20

CD_3_OD

28

29

15

13

1

25

23

27

11

3

11

3

27

Figure S16. ^1^H-NMR of Accramycin A **1** (CD_3_OD, 298K, 600MHz)

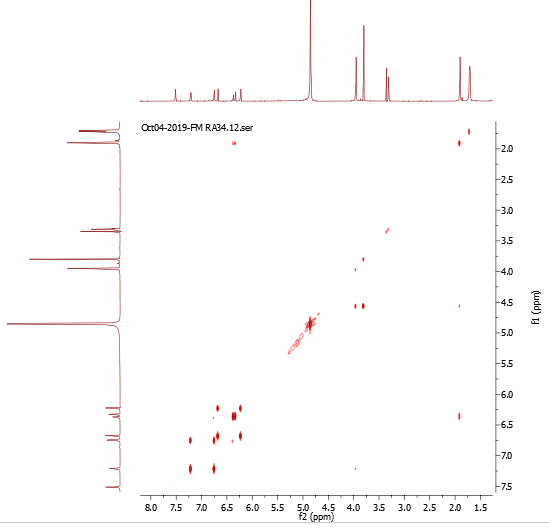

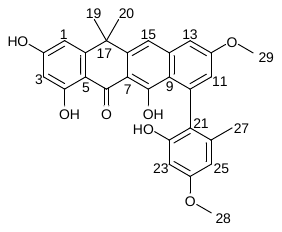

19, 20

19, 20

27

27

CD_3_OD

28

29

29

CD_3_OD

28

23

3

25

11

1

15

13

15

13

11

3

1

23

25

25

28

1

Figure S17. ^1^H-^1^H COSY NMR of Accramycin A **1** (CD_3_OD, 298K, 600MHz)

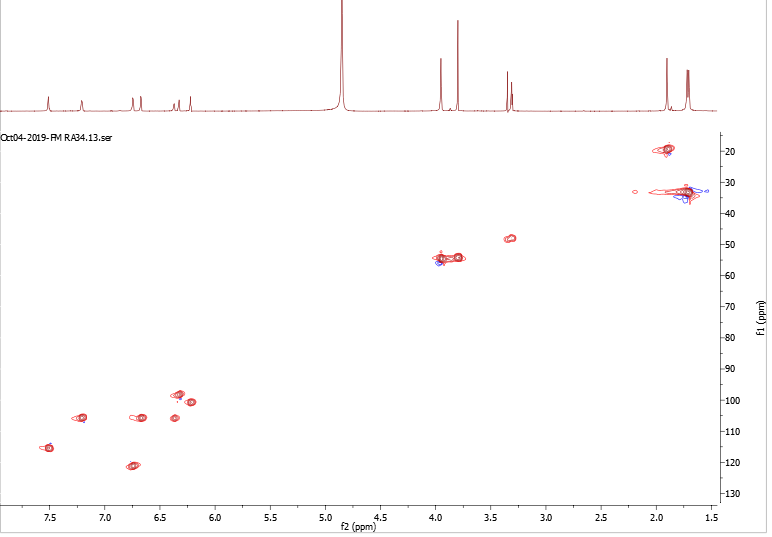

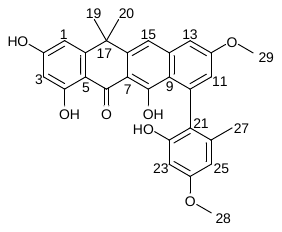

19, 20

27

28

29

3

15

13

11

1

25

23

MeOD

Figure S18. HSQC of Accramycin A **1** (CD_3_OD, 298K, 600MHz)

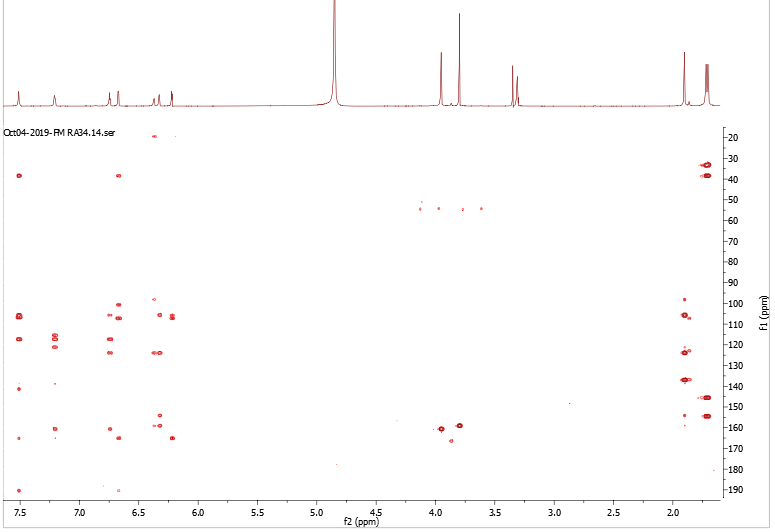

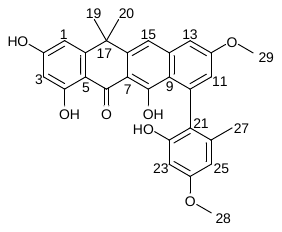

19, 20

27

CD_3_OD

28

29

3

23

25

1

11

13

15

Figure S19. HMBC of Accramycin A **1** (CD_3_OD, 298K, 600MHz)

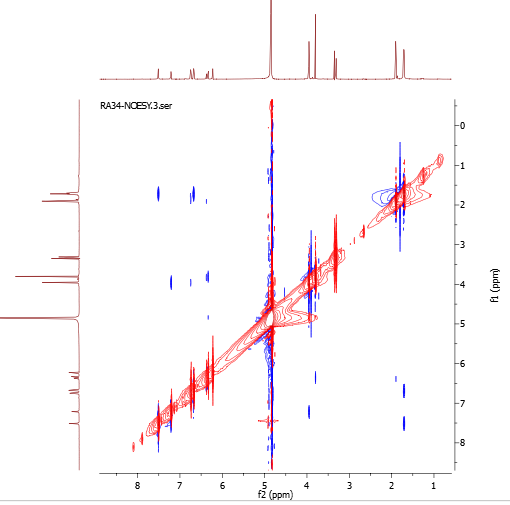

19, 20

19, 20

27

27

29

28

28

29

25, 23, 3

11

1

15

13

15

13

11, 1

25, 23

3

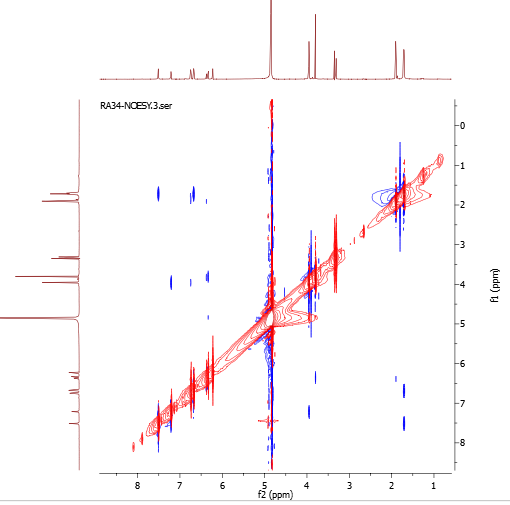

19, 20

19, 20

Figure S20. NOESY of Accramycin A **1** (CD_3_OD, 298K, 600MHz)

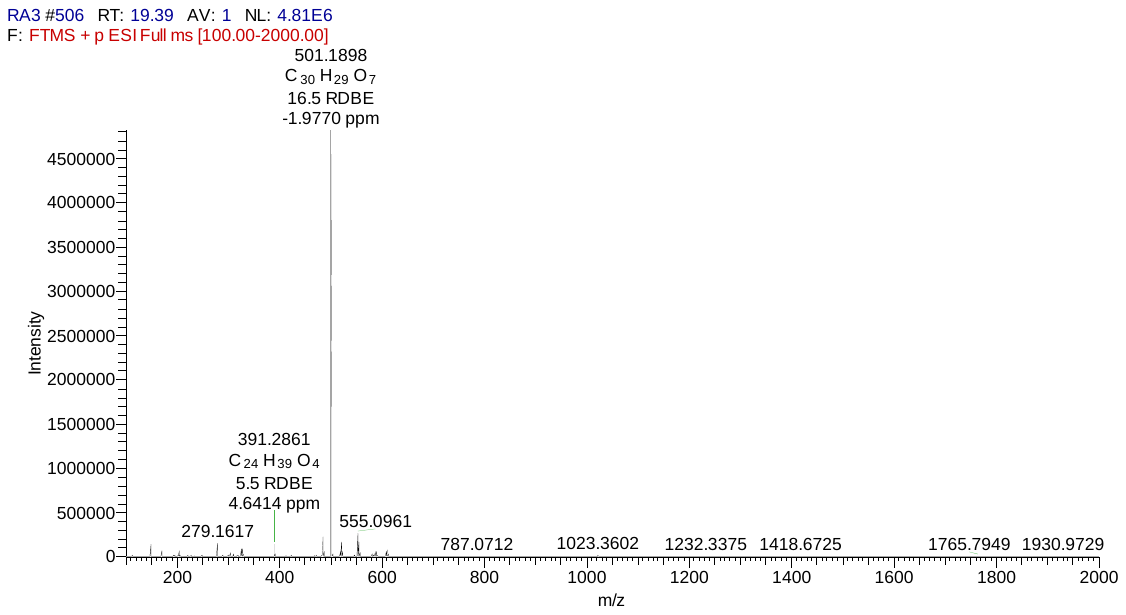

[M+H]^+^

Figure S21. HRESIMS of Accramycin B **2**

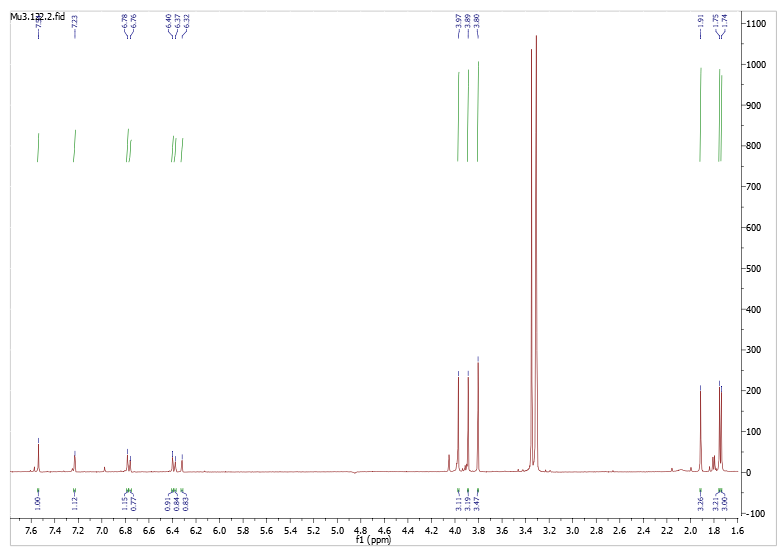

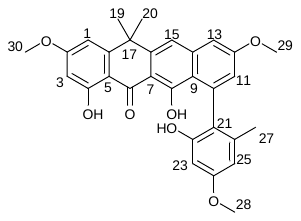

19, 20

27

28

29

30

15

13

11 1

25, 23, 3

Figure S22. ^1^H-NMR of Accramycin B **2** (CD_3_OD, 298K, 600MHz)

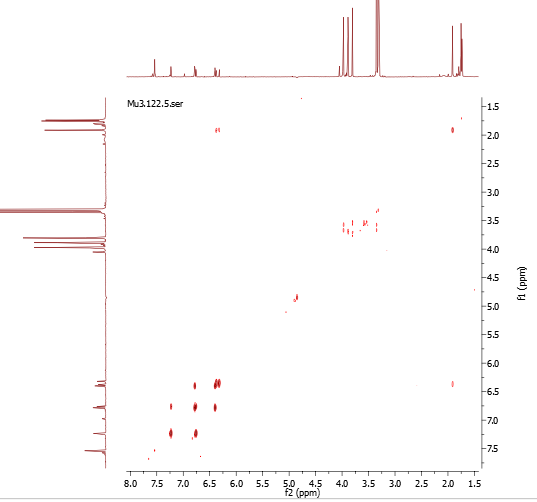

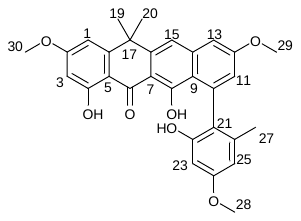

Figure S23. COSY of Accramycin B **2** (CD_3_OD, 298K, 600MHz)

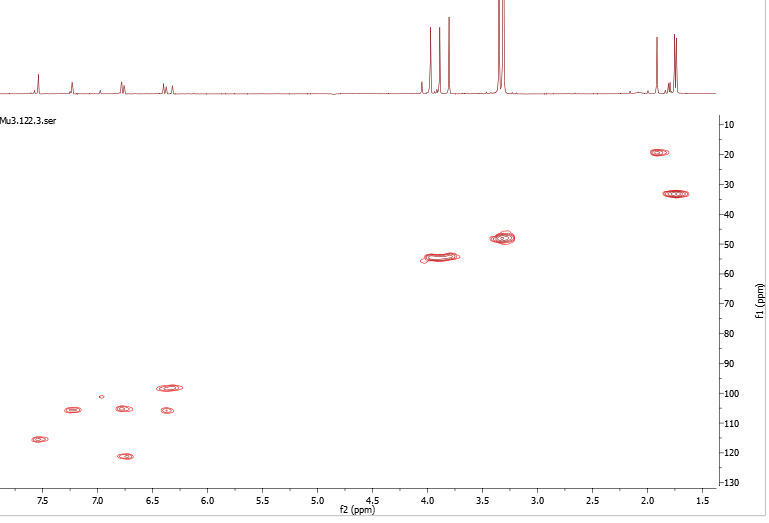

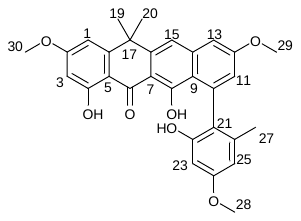

Figure S24. HSQC of Accramycin B **2** (CD_3_OD, 298K, 600MHz)

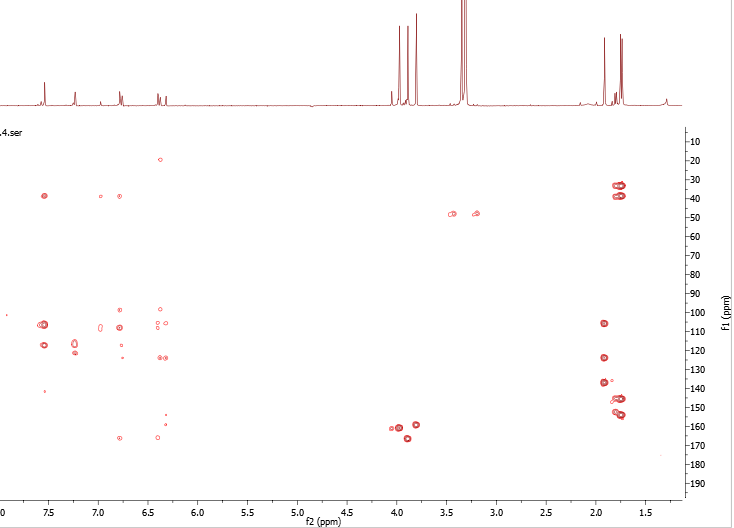

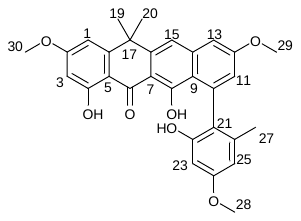

Figure S25. HMBC of Accramycin B **2** (CD_3_OD, 298K, 600MHz)

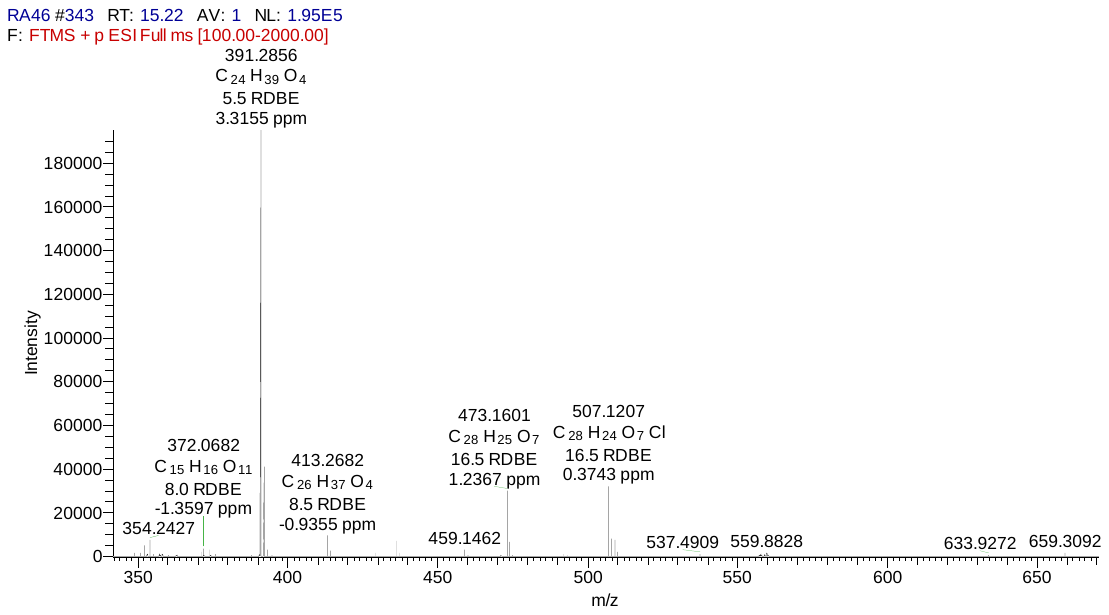

[M+H]^+^

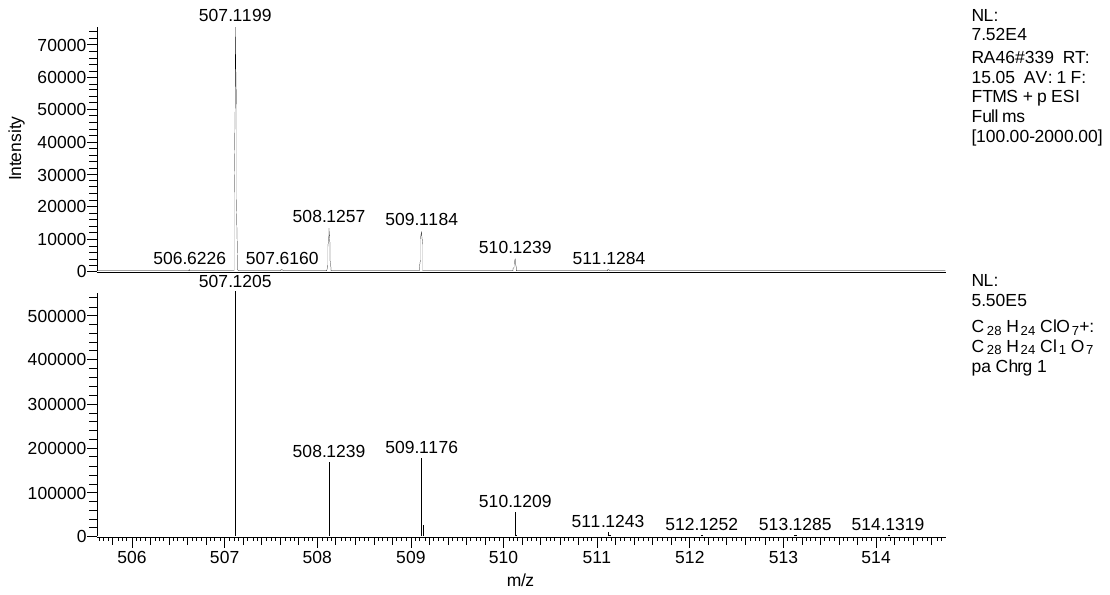

calculated

observed

**A**

**B**

Figure S26. **A**. HRESIMS and **B.** Isotope Pattern of Accramycin C **3**

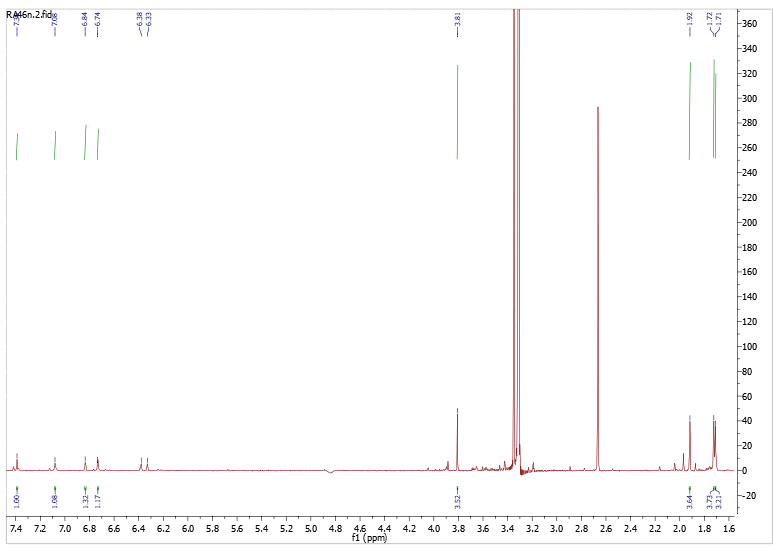

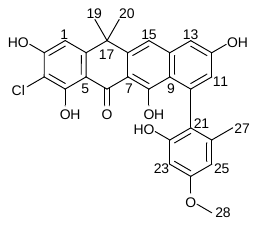

19, 20

27

28

15

13

11

1

25

23

Figure S27. ^1^H-NMR of Accramycin C **3** (CD_3_OD, 298K, 600MHz)

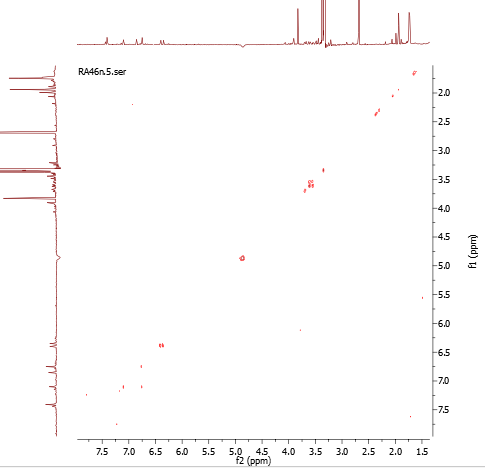

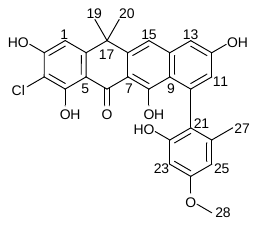

25, 23

25, 23

13

11

11

13

19, 20

27

28

15

1

19, 20

27

28

15

1

Figure S28. ^1^H-^1^H COSY of Accramycin C **3** (CD_3_OD, 298K, 600MHz)

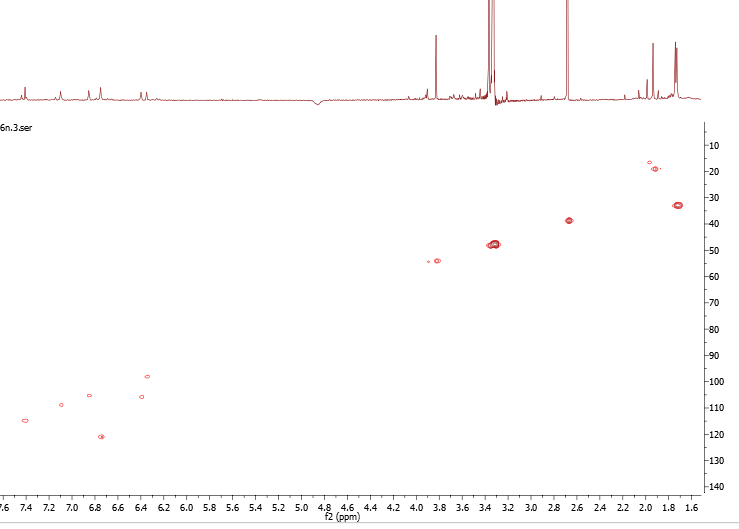

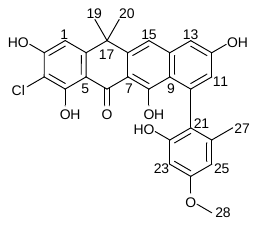

19, 20

27

28

15

11

13

1

25

23

Figure S29. HSQC of Accramycin C **3** (CD_3_OD, 298K, 600MHz)

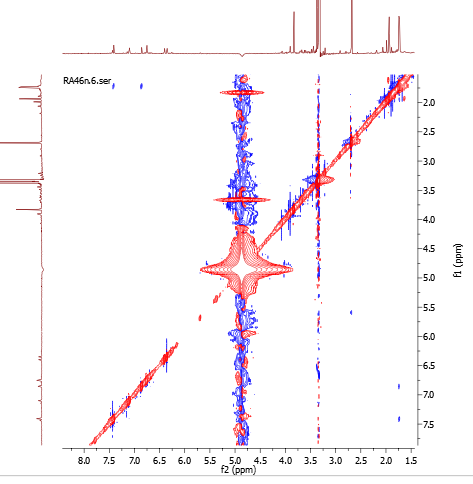

19, 20

15

1

19, 20

27

27

28

28

13

25, 23

11

25, 23

15

13

1

11

Figure S30. NOESY of Accramycin C **3** (CD_3_OD, 298K, 600MHz)

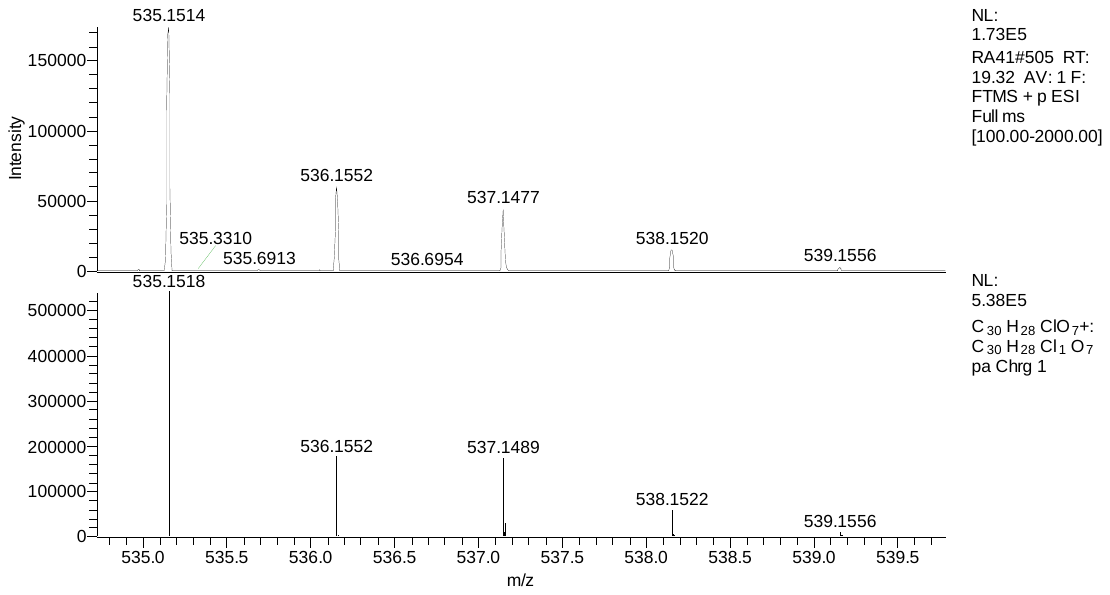

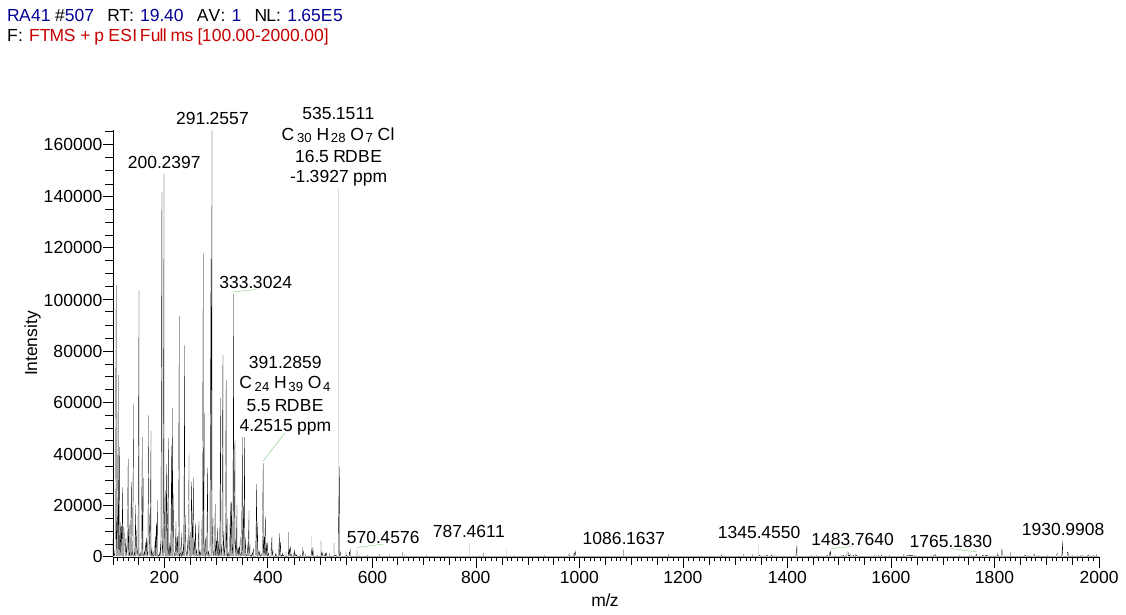

[M+H]^+^

observed

calculated

**A**

**Bcc**

Figure S31. **A**. HRESIMS and **B**. Isotope Pattern of Accramycin D **4**

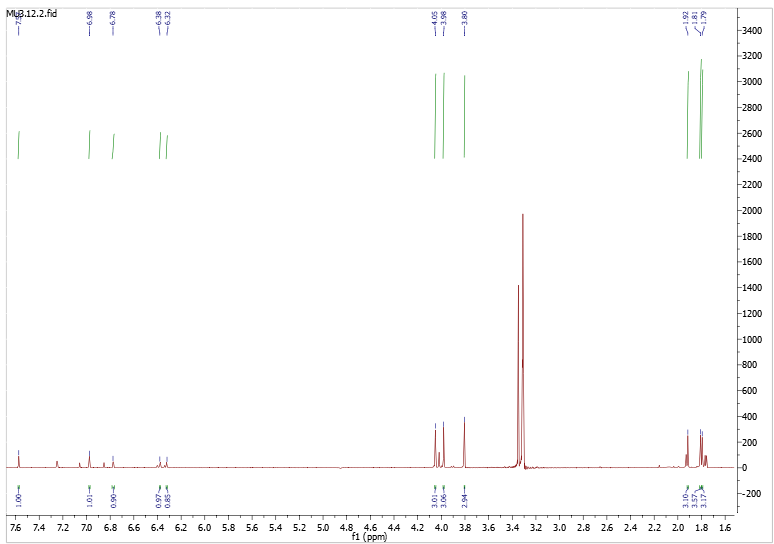

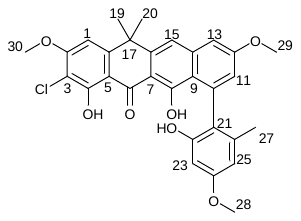

Figure S32. ^1^H-NMR of Accramycin D **4** (CD_3_OD, 298K, 600MHz)

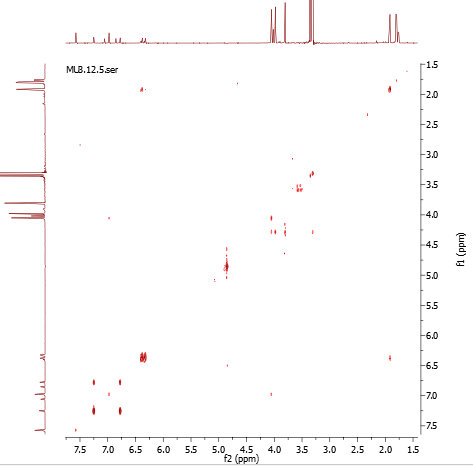

Figure S33. ^1^H-^1^H COSY of Accramycin D **4** (CD_3_OD, 298K, 600MHz)

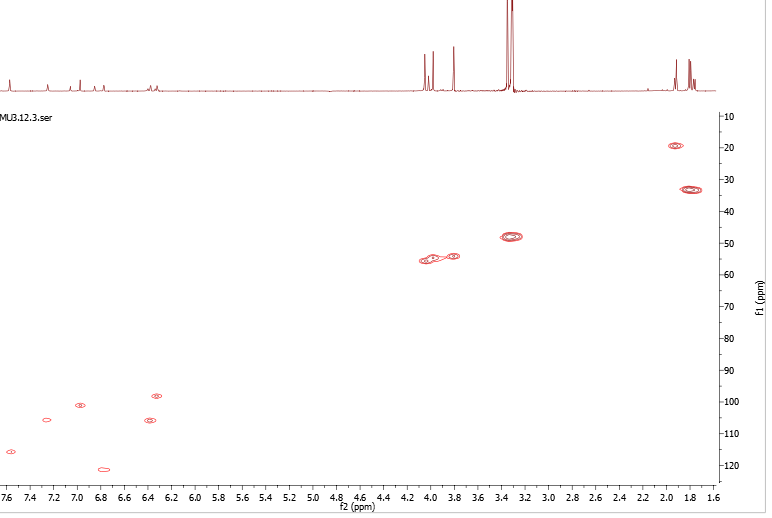

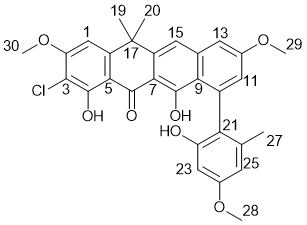

Figure S34. HSQC of Accramycin D **4** (CD_3_OD, 298K, 600MHz)

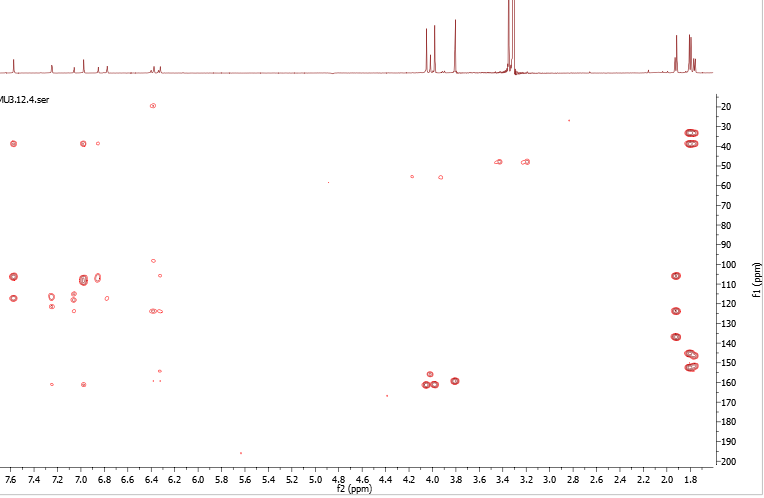

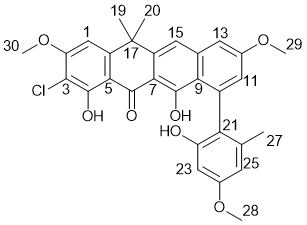

Figure S35. HMBC of Accramycin D **4** (CD_3_OD, 298K, 600MHz)

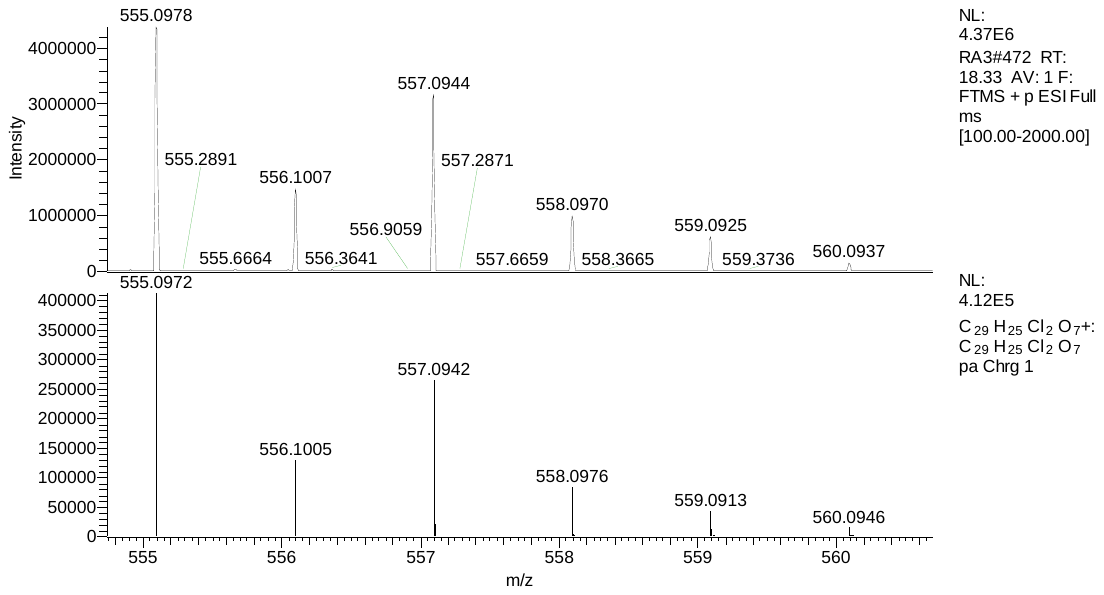

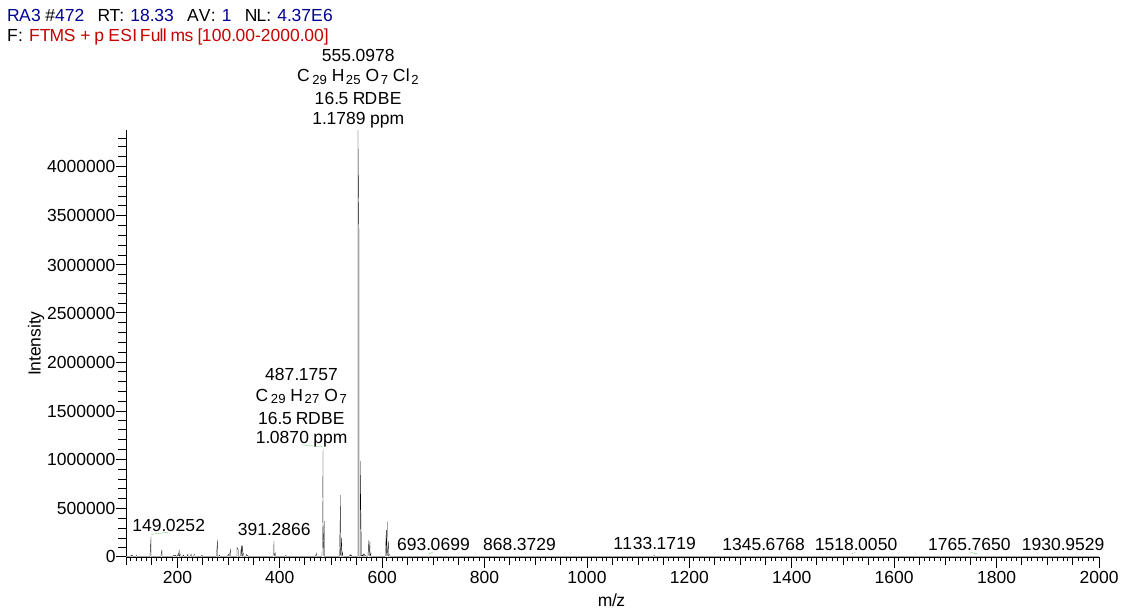

[M+H]^+^

observed

calculated

**A**

**B**

Figure S36. **A.** HRESIMS and **B.** Isotope Pattern of Accramycin E **5**

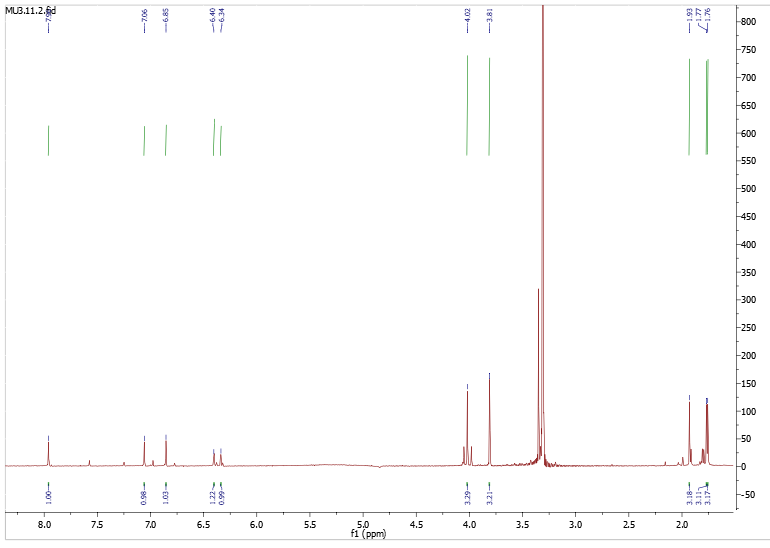

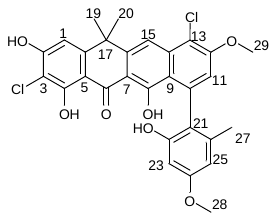

19, 20

27

28

29

15

25, 23

11

1

Figure S37. ^1^H-NMR Accramycin E **5** (CD_3_OD, 298K, 600MHz)

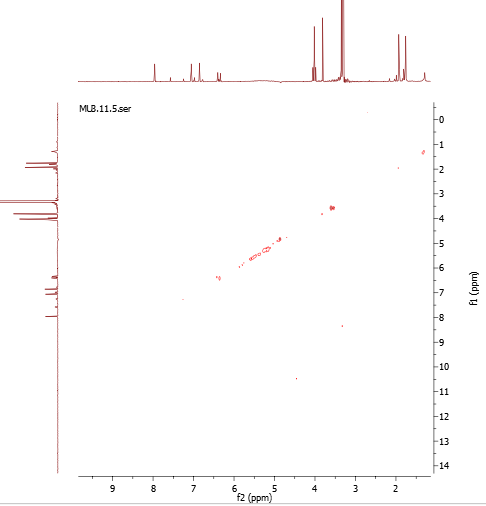

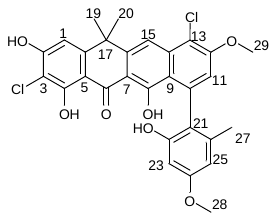

Figure S38. ^1^H-^1^H COSY Accramycin E **5** (CD_3_OD, 298K, 600MHz)

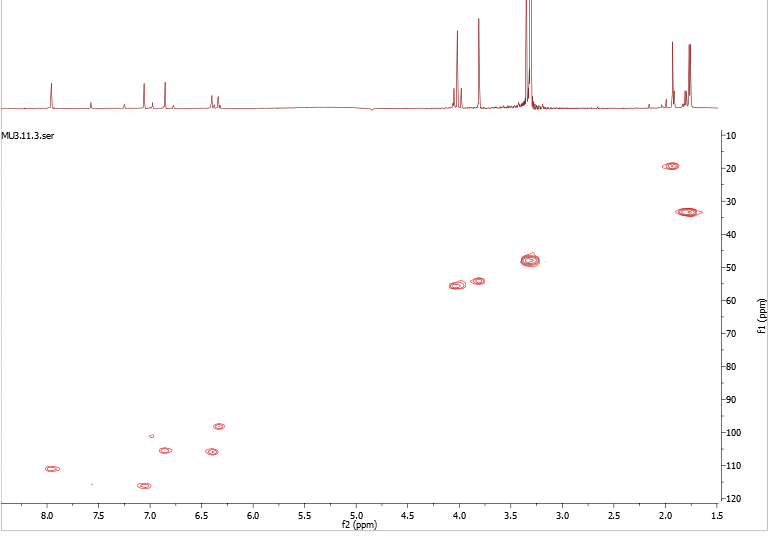

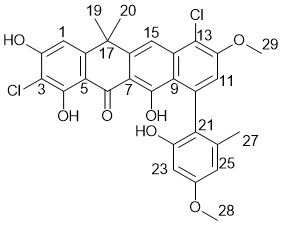

15

11

1

25

23

29

28

27

19, 20

Figure S39. HSQC Accramycin E **5** (CD_3_OD, 298K, 600MHz)

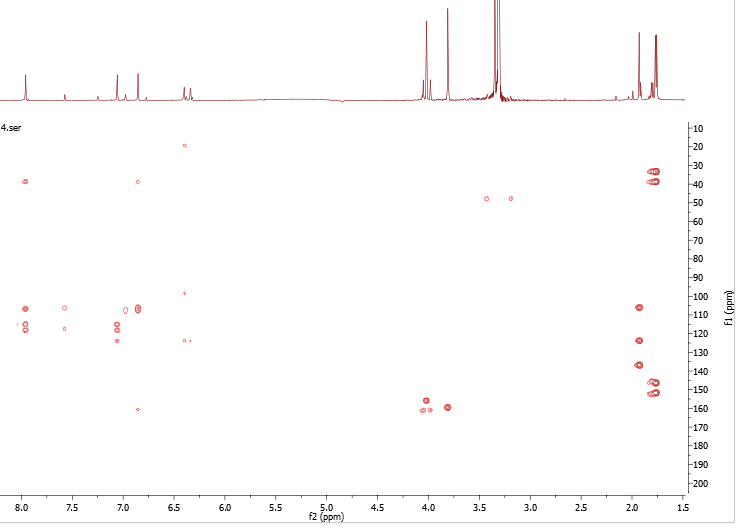

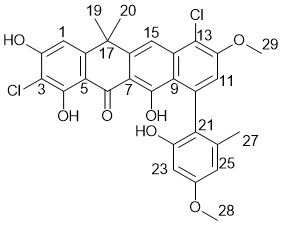

15

11

1

25

23

29

28

27

19, 20

Figure S40. HMBC Accramycin E **5** (CD_3_OD, 298K, 600MHz)

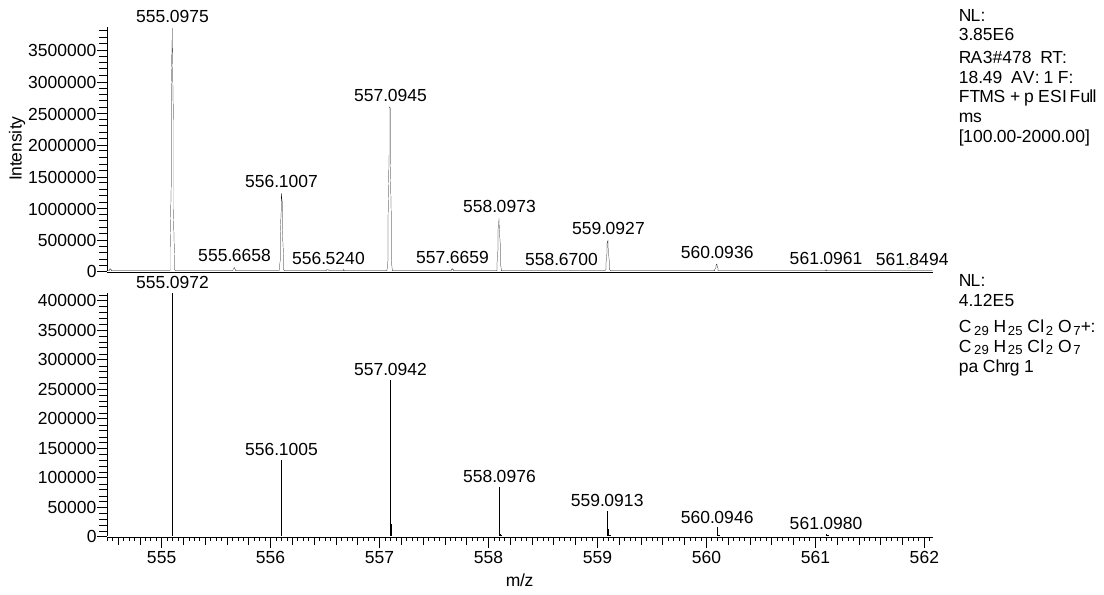

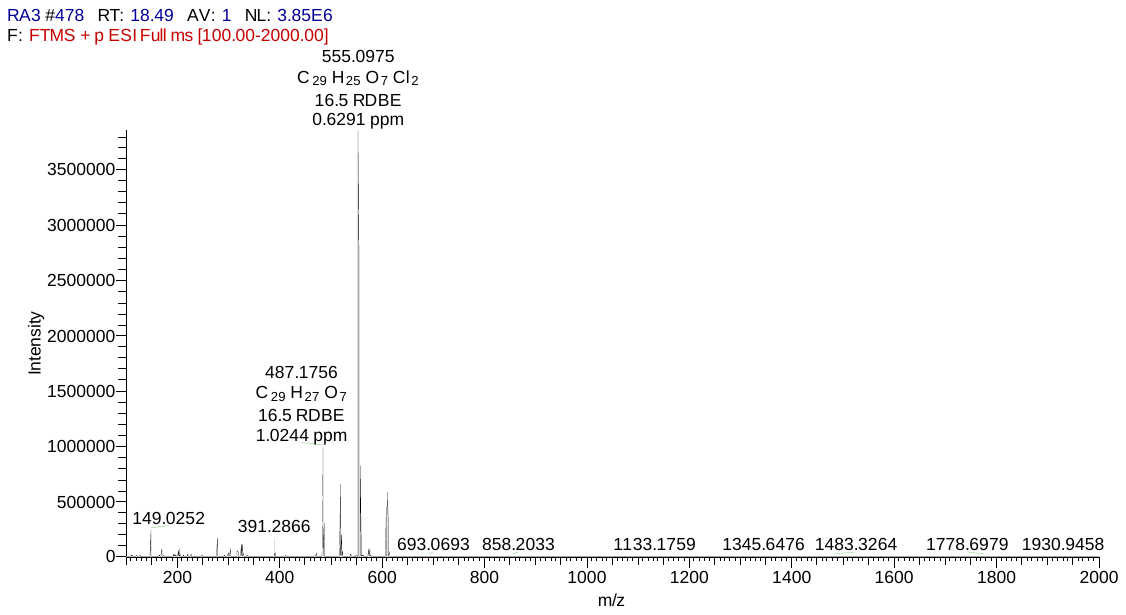

[M+H]^+^

calculated

observed

**A**

**B**

Figure S41. **A.** HRESIMS and **B.** Isotope Pattern of Accramycin F **6**

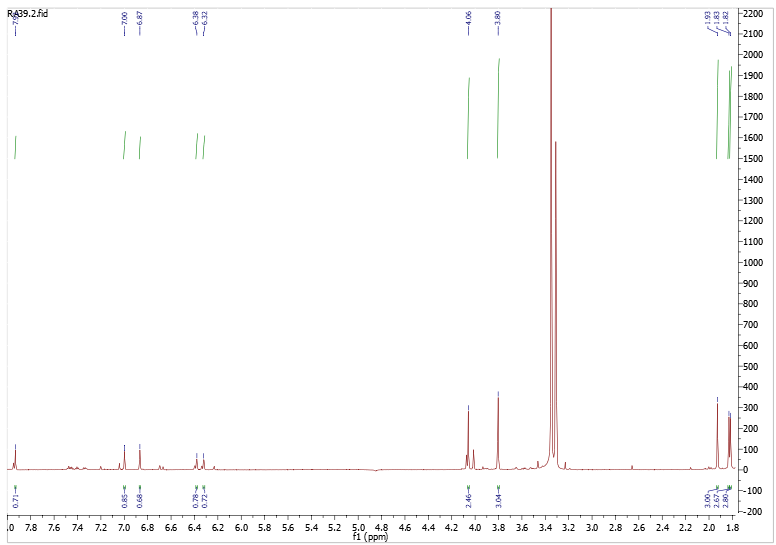

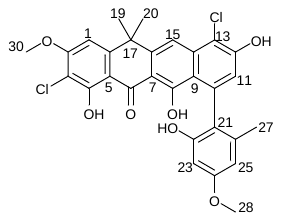

27

19, 20

28

30

15

25

23

1

11

Figure S42. ^1^H-NMR of Accramycin F **6** (CD_3_OD, 298K, 600MHz)

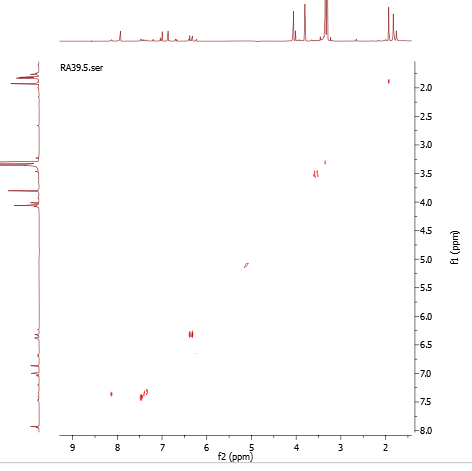

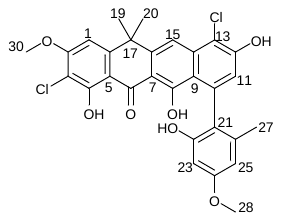

23

25

23

25

15

1

11

30

28

27

19, 20

19, 20

27

28

30

15

1

11

Figure S43. ^1^H-^1^H COSY of Accramycin F **6** (CD_3_OD, 298K, 600MHz)

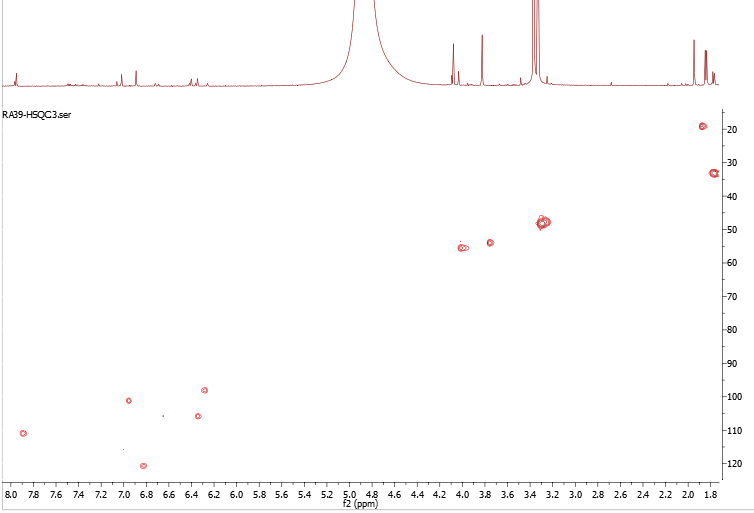

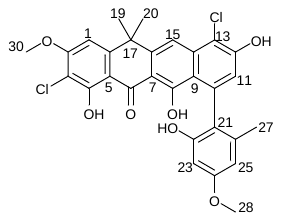

15

1

11

25

23

30

28

27

19, 20

Figure S44. HSQC NMR of Accramycin F **6** (CD_3_OD, 298K, 600MHz)

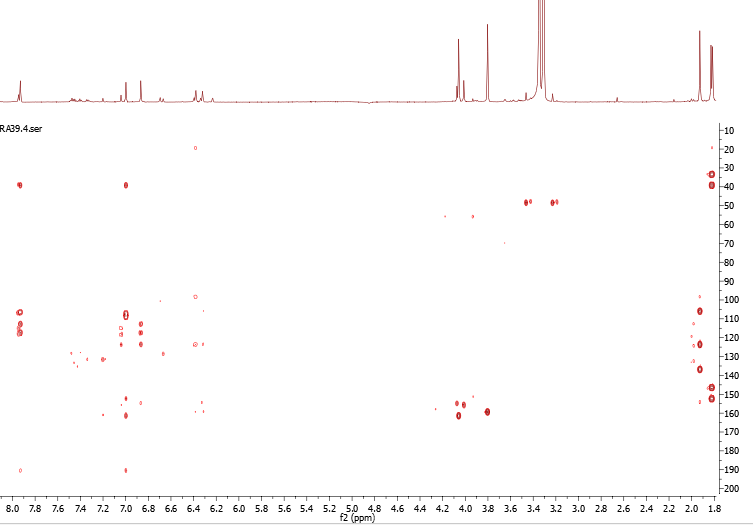

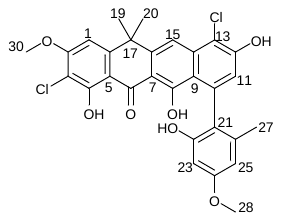

15

1

11

25

23

30

28

27

19, 20

Figure S45. HMBC NMR of Accramycin F **6** (CD_3_OD, 298K, 600MHz)

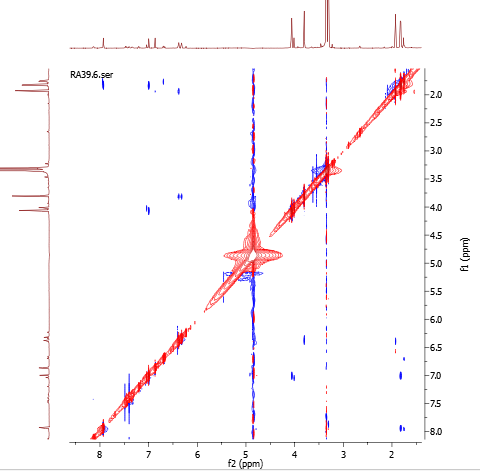

15

1

11

25

23

30

28

27

19, 20

19, 20

27

28

30

23

25

11

1

15

Figure S46. NOESY of Accramycin F **6** (CD_3_OD, 298K, 600MHz)

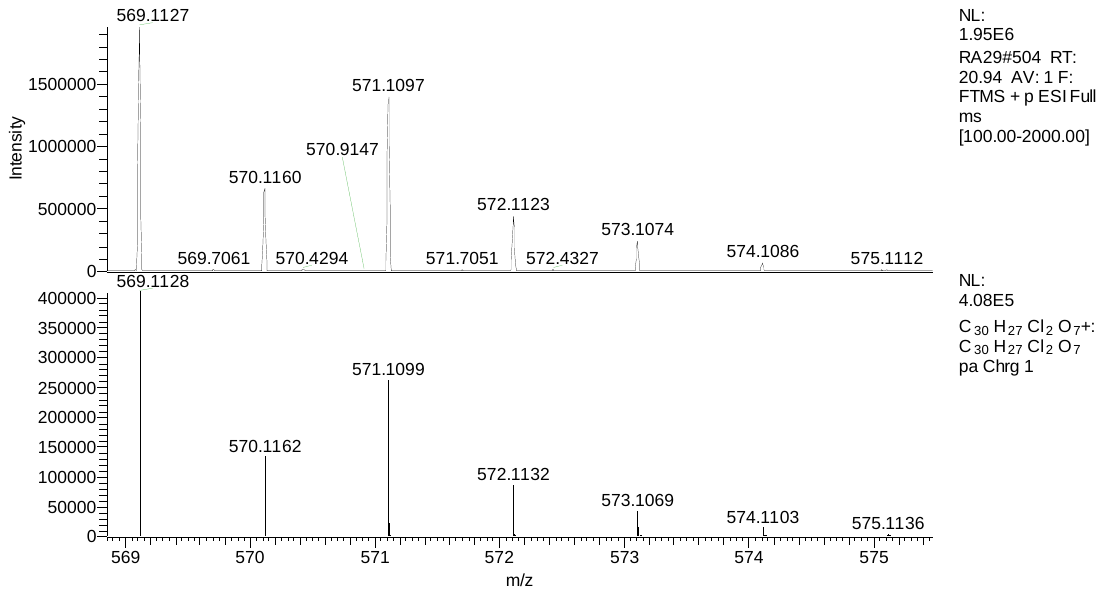

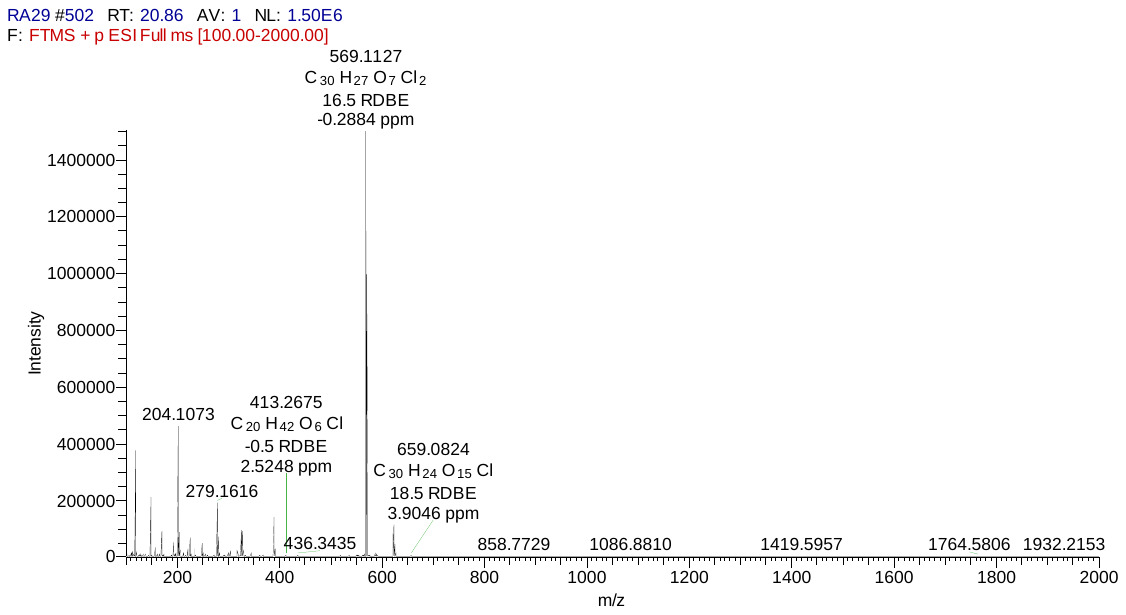

[M+H]^+^

calculated

observed

**A**

**B**

Figure S47. **A.** HRESIMS and **B.** Isotope Pattern of Accramycin G **7**

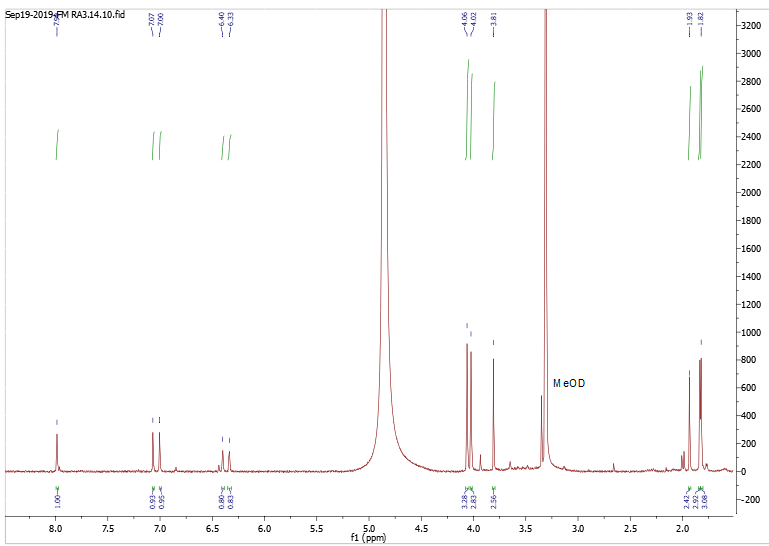

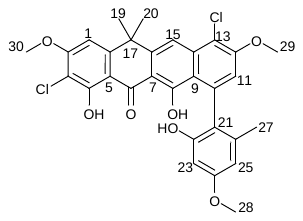

19, 20

27

28

29

30

15

11

1

25

23

Figure S48. ^1^H-NMR of Accramycin G **7** (CD_3_OD, 298K, 600MHz)

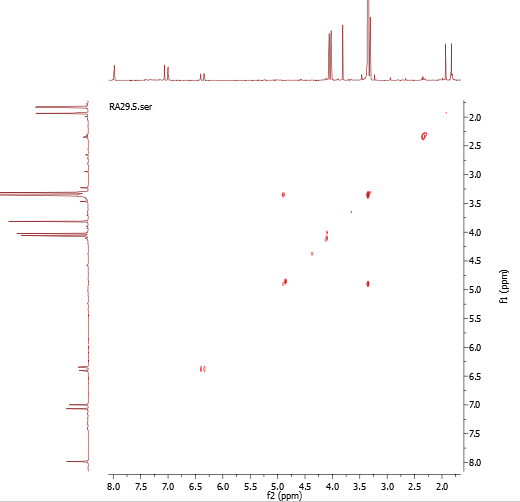

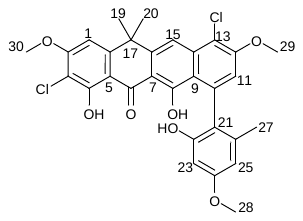

25

23

25

23

27

19, 20

19, 20

27

30

29

28

15

11

1

30

29

28

15

11

1

Figure S49. ^1^H-^1^H COSY of Accramycin G **7** (CD_3_OD, 298K, 600MHz)

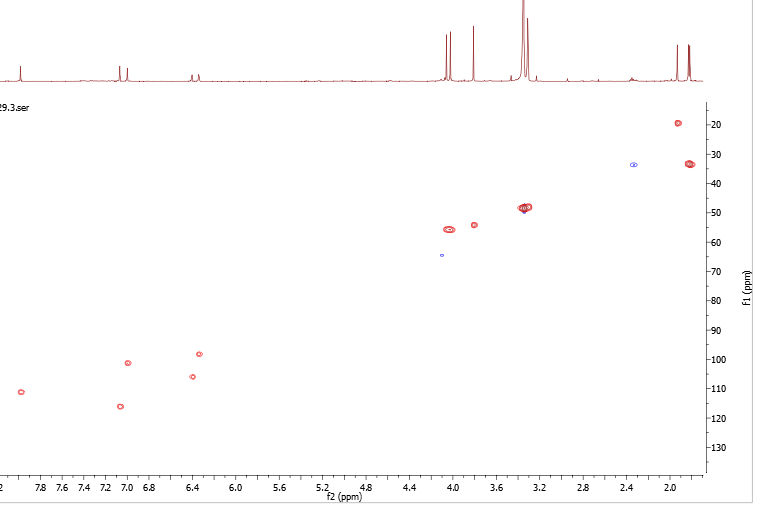

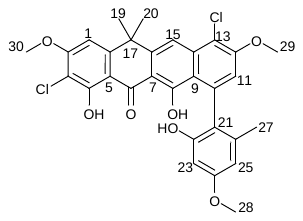

15

11

1

25

23

30

29

28

27

19, 20

Figure S50. HSQC of Accramycin G **7** (CD_3_OD, 298K, 600MHz)

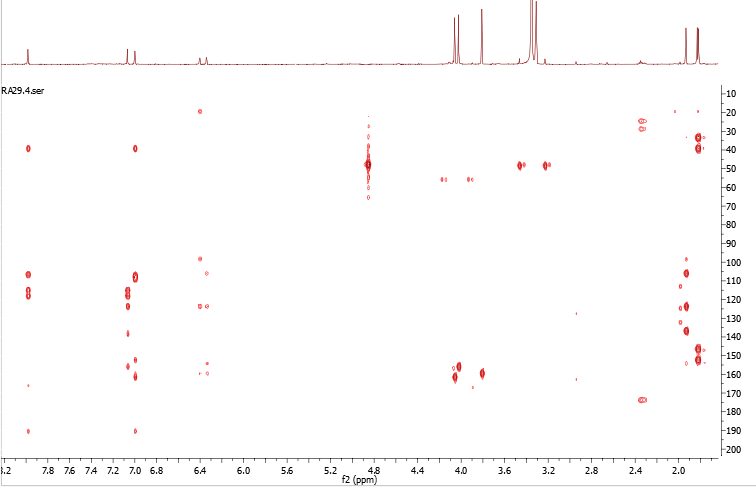

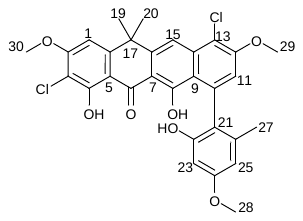

15

11

1

25

23

30

29

28

27

19, 20

Figure S51. HMBC of Accramycin G **7** (CD_3_OD, 298K, 600MHz)

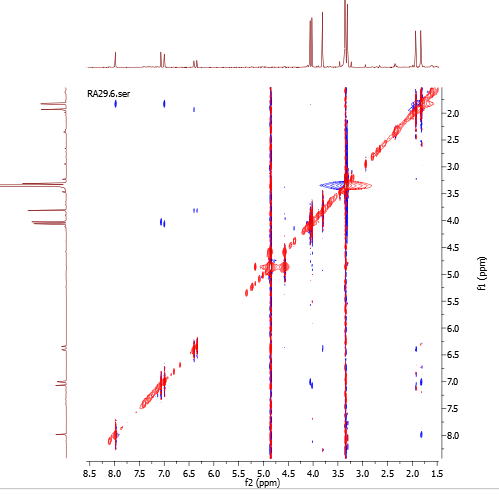

15

11

1

25

23

30

29

28

27

19, 20

19, 20

27

28

30

29

25

23

1

11

15

Figure S52. NOESY of Accramycin G **7** (CD_3_OD, 298K, 600MHz)

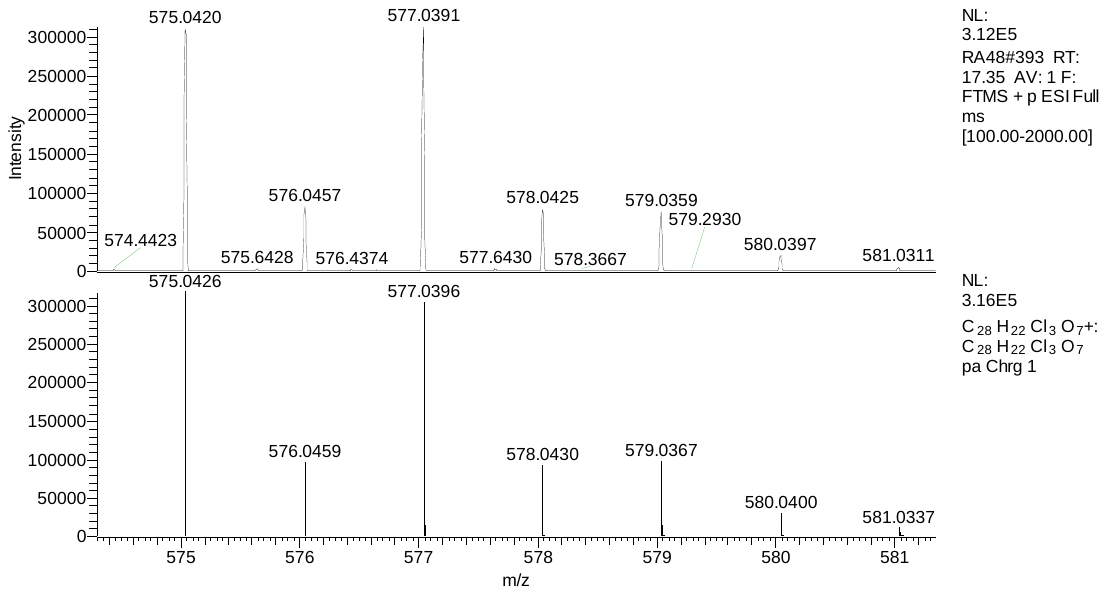

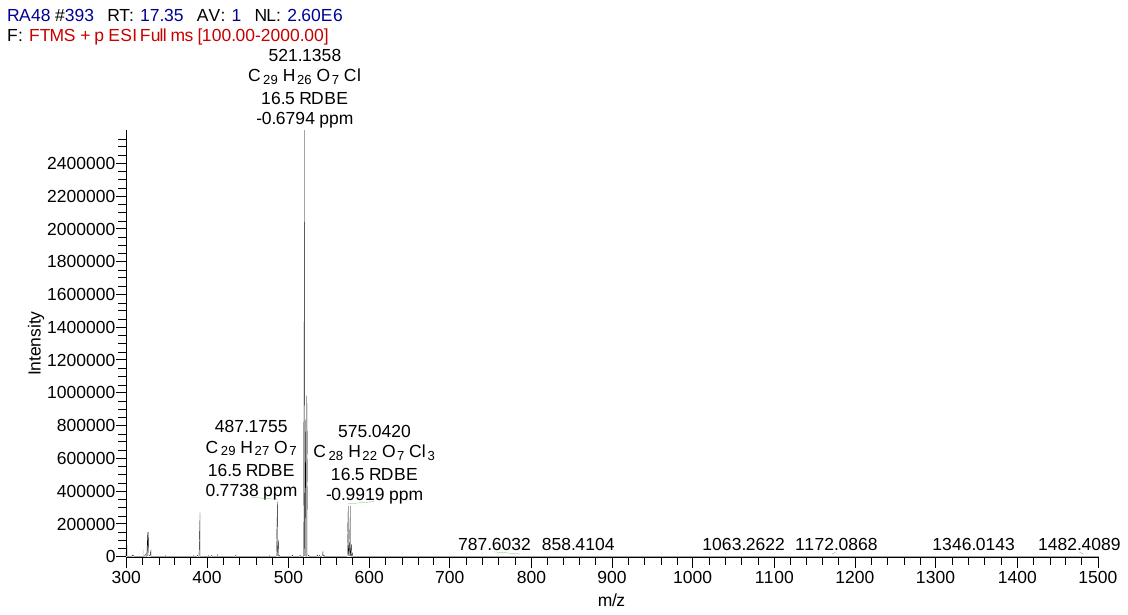

[M+H]^+^

calculated

observed

**A**

**B**

Figure S53. **A.** HRESIMS and **B.** Isotope Pattern of Accramycin H **8**

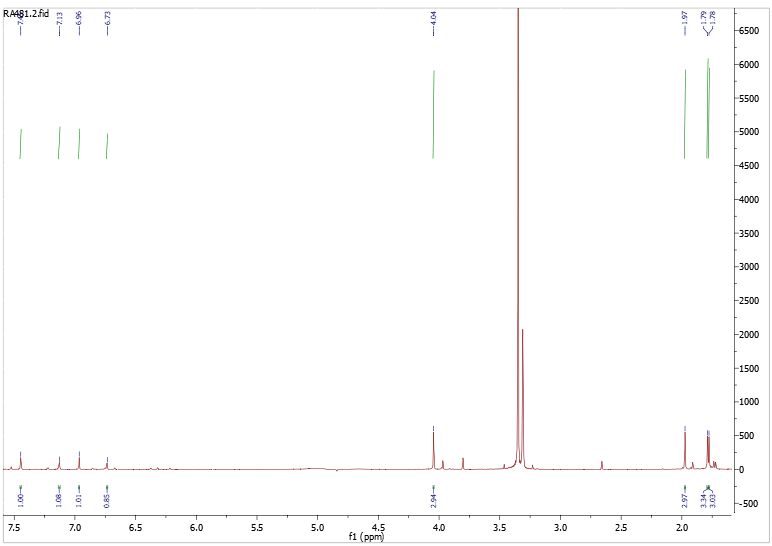

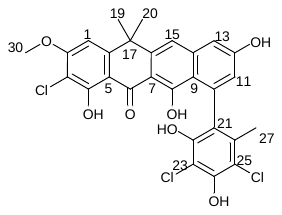

19, 20

27

30

15

13

1

11

Figure S54. ^1^H-NMR of Accramycin H **8** (CD_3_OD, 298K, 600MHz)

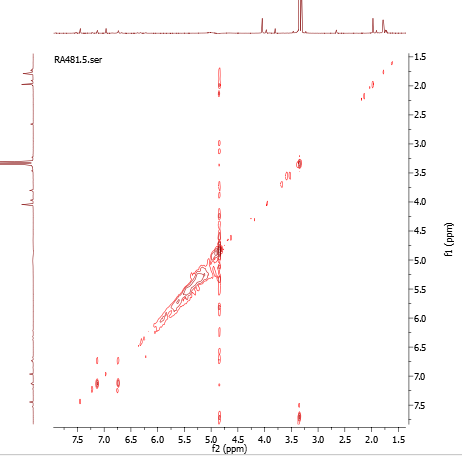

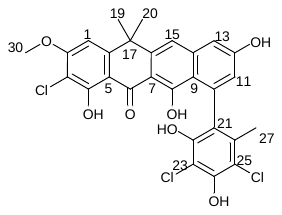

15

1

11

13

30

27

19, 20

30

27

19, 20

Figure S55. ^1^H-^1^H COSY of Accramycin H **8** (CD_3_OD, 298K, 600MHz)

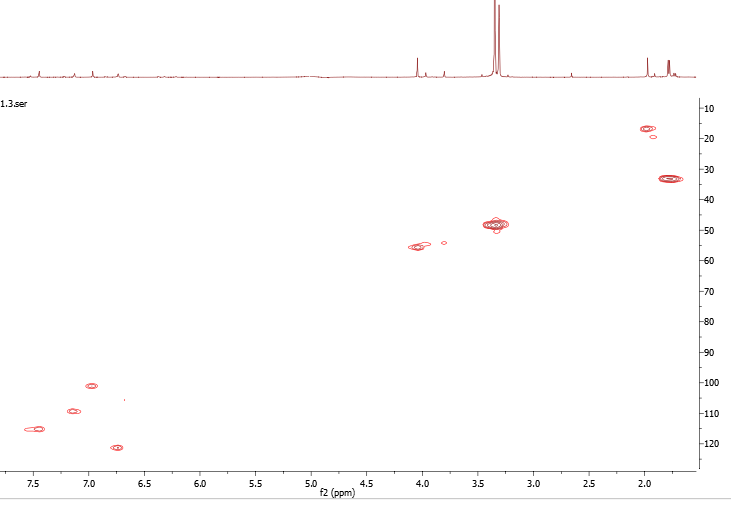

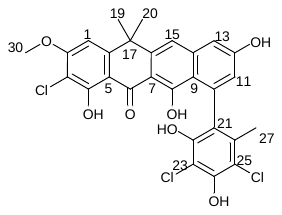

15

13

1

11

30

27

19, 20

Figure S56. HSQC of Accramycin H **8** (CD_3_OD, 298K, 600MHz)

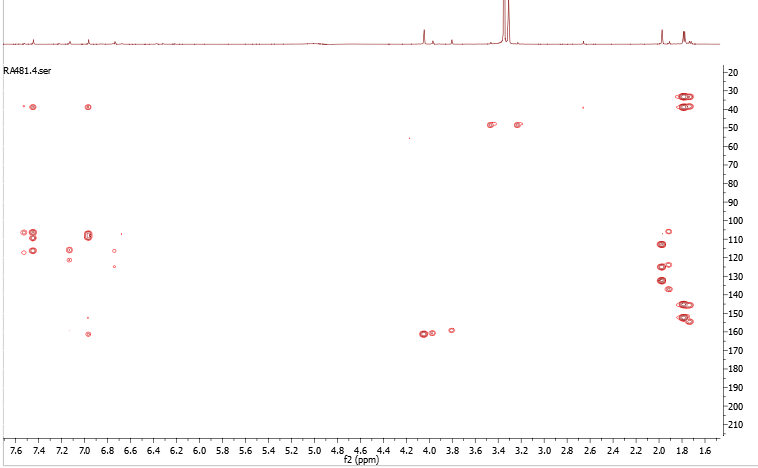

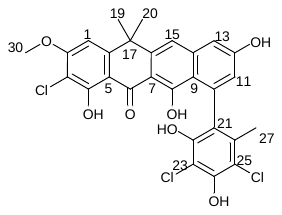

15

13

1

11

30

27

19, 20

Figure S57. HMBC of Accramycin H **8** (CD_3_OD, 298K, 600MHz)

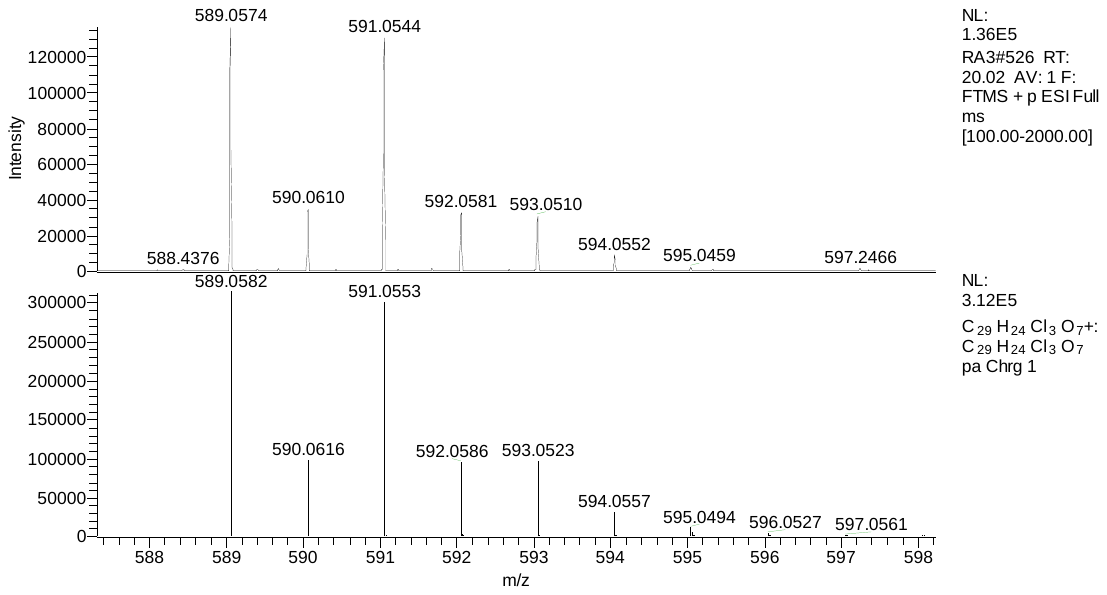

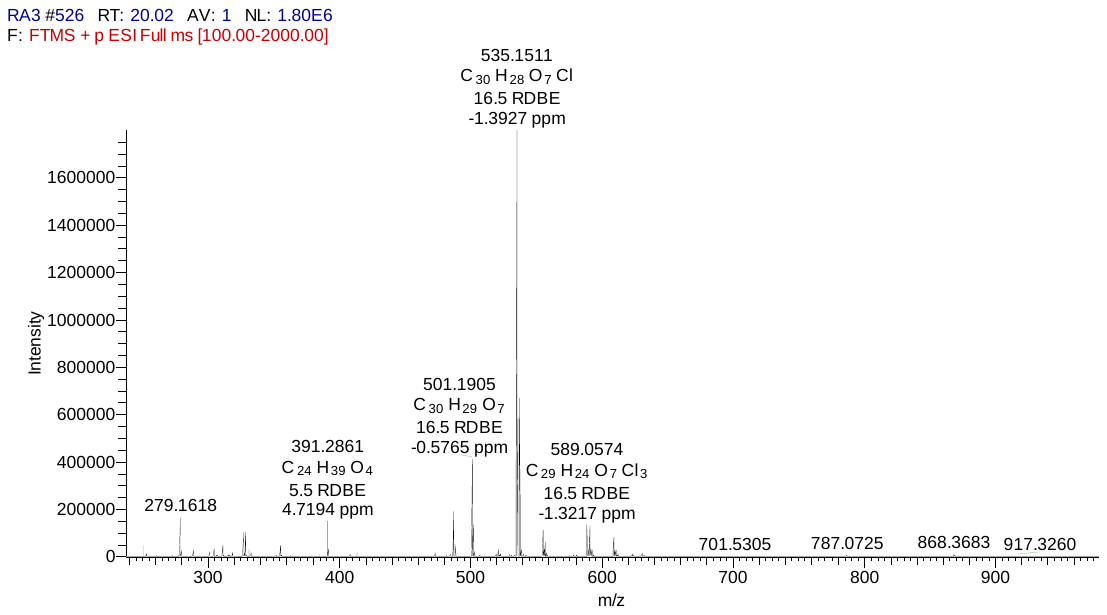

[M+H]^+^

calculated

observed

**A**

**B**

Figure S58. **A.** HRESIMS and **B.** Isotope Pattern of Accramycin I **9**

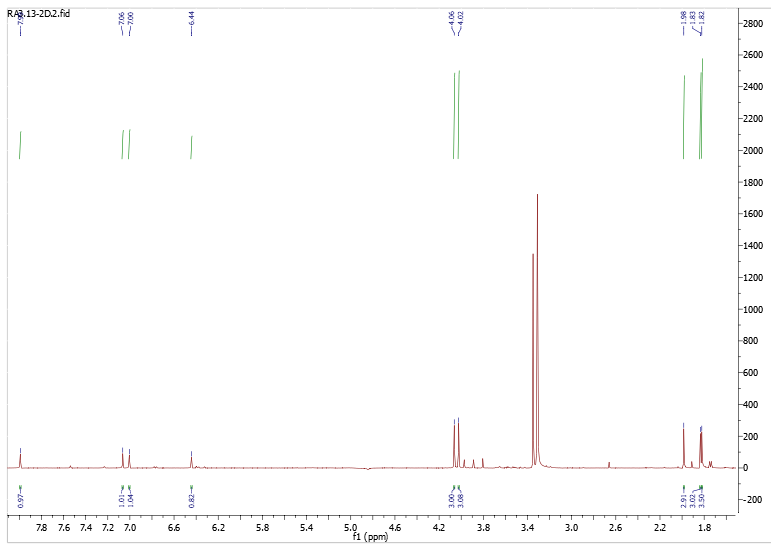

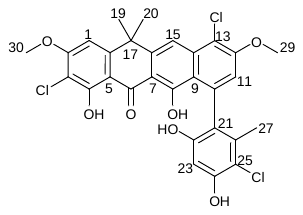

19, 20

27

29

30

23

15

11

1

Figure S59. ^1^H NMR of Accramycin I **9** (CD_3_OD, 298K, 600MHz)

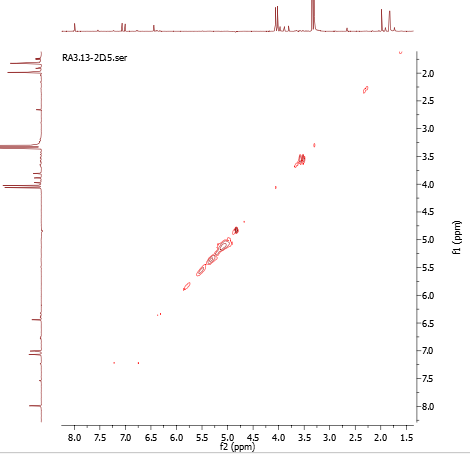

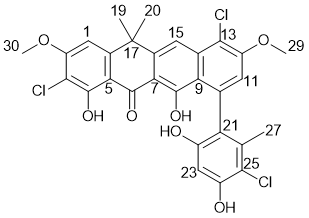

Figure S60. ^1^H-^1^H COSY of Accramycin I **9** (CD_3_OD, 298K, 600MHz)

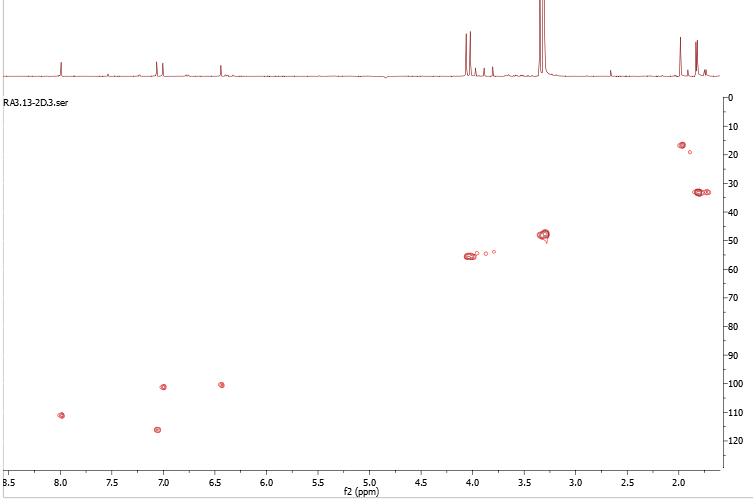

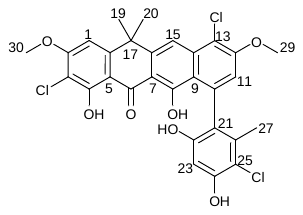

15

11

1

23

30

29

27

19, 20

Figure S61. HSQC of Accramycin I **9** (CD_3_OD, 298K, 600MHz)

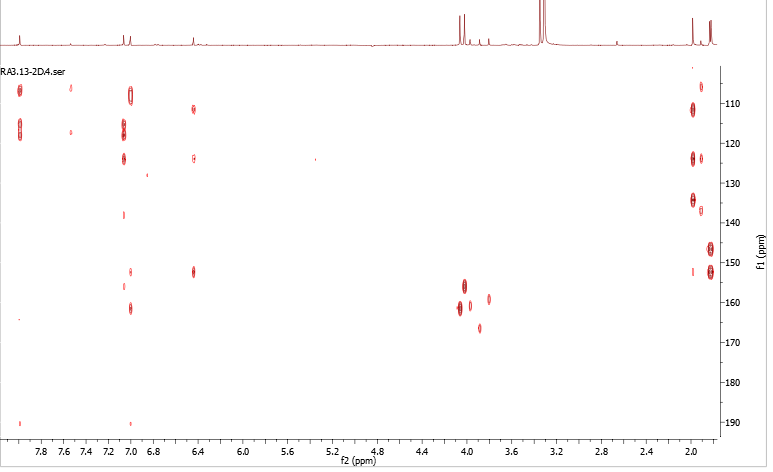

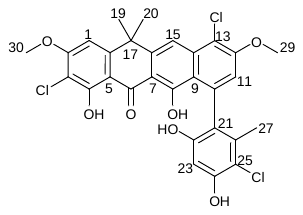

15

11

1

23

30

29

27

19, 20

Figure S62. HMBC of Accramycin I **9** (CD_3_OD, 298K, 600MHz)

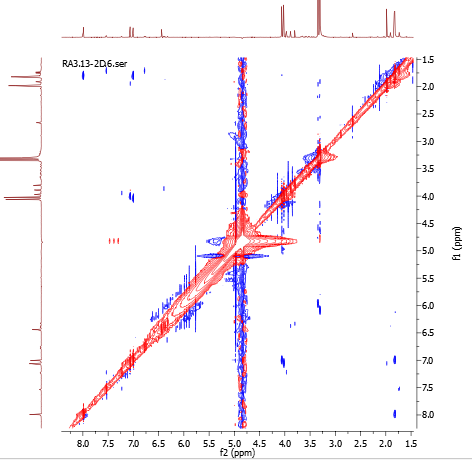

15

23

11

1

19, 20

27

30

29

27

19, 20

29

30

23

11

1

15

Figure S63. NOESY of Accramycin I **9** (CD_3_OD, 298K, 600MHz)

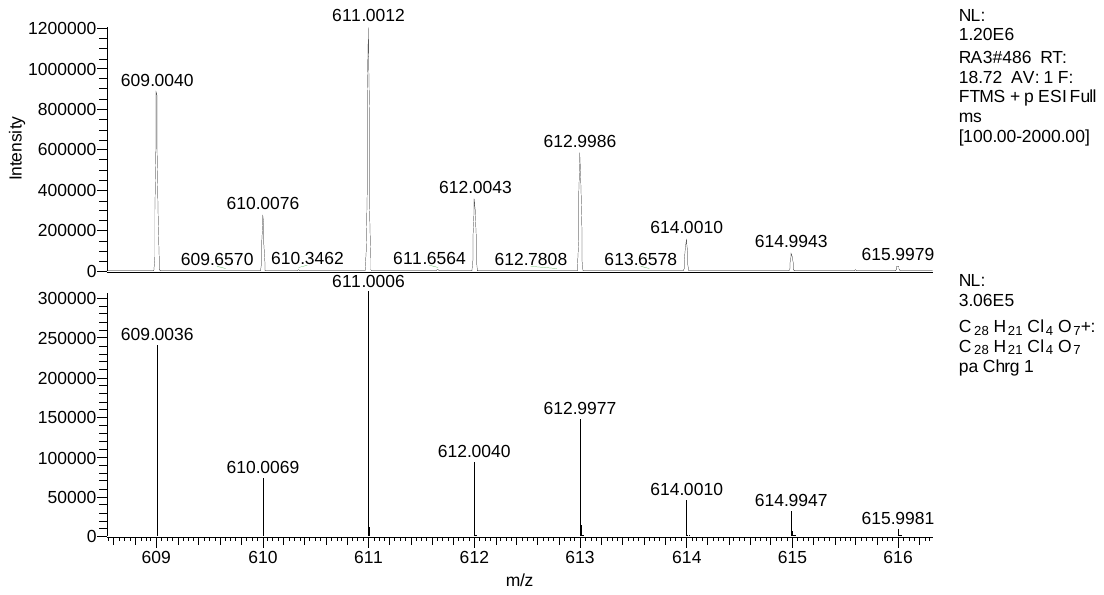

calculated

observed

Figure S64. LCMS isotope pattern of Accramycin J **10**

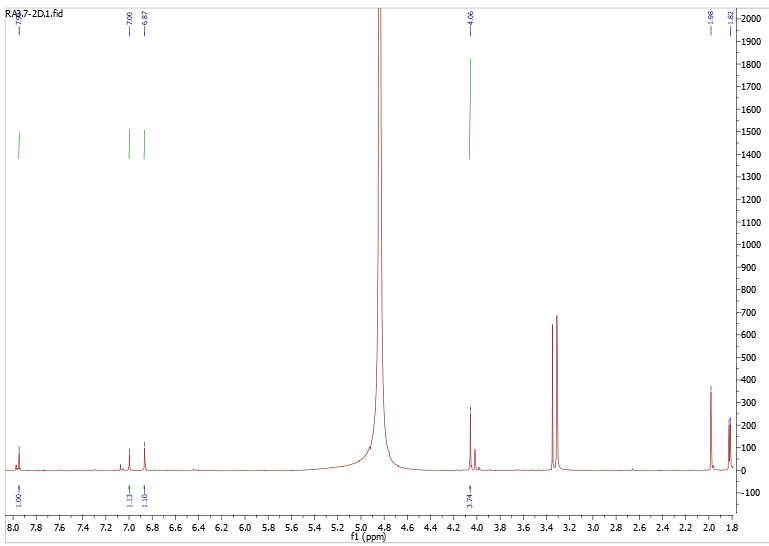

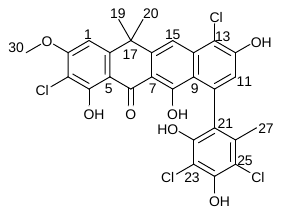

15

1

11

30

27

19,20

Figure S65. ^1^H-NMR of Accramycin J **10** (CD_3_OD, 298K, 600MHz)

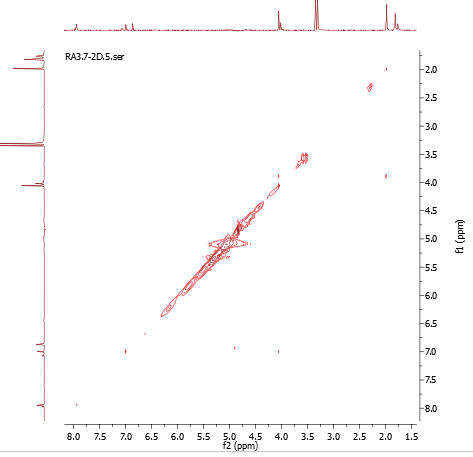

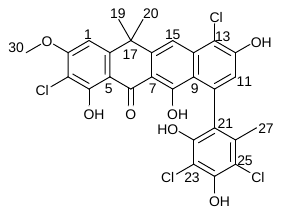

15

15

1

11

30

27

19, 20

27

19, 20

30

1

11

Figure S66. ^1^H-^1^H COSY of Accramycin J **10** (CD_3_OD, 298K, 600MHz)

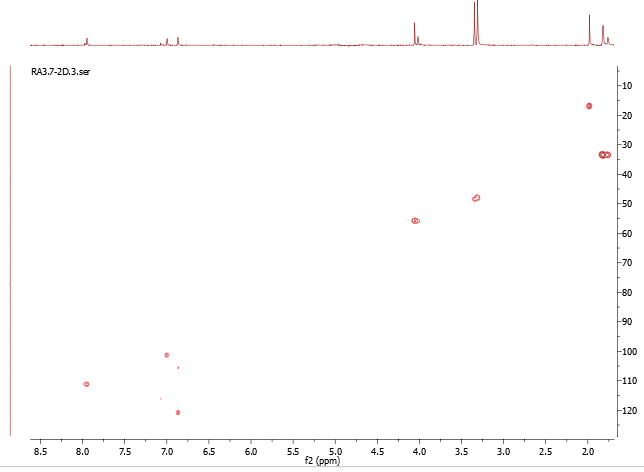

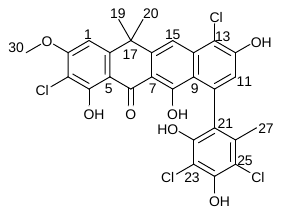

15

1

11

30

27

19, 20

Figure S67. HSQC of Accramycin J **10** (CD_3_OD, 298K, 600MHz)

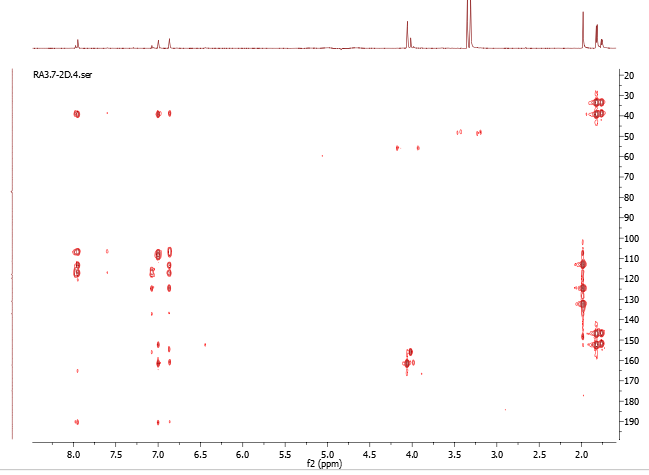

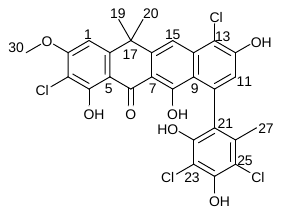

15

1

11

30

27

19, 20

Figure S68. HMBC of Accramycin J **10** (CD_3_OD, 298K, 600MHz)

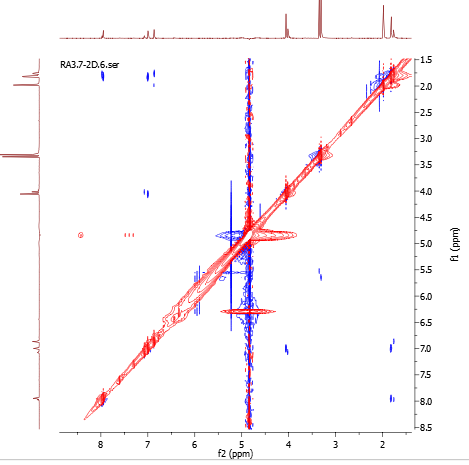

15

1

11

30

30

19, 20

27

27

19, 20

Figure S69. NOESY of Accramycin J **10** (CD_3_OD, 298K, 600MHz)

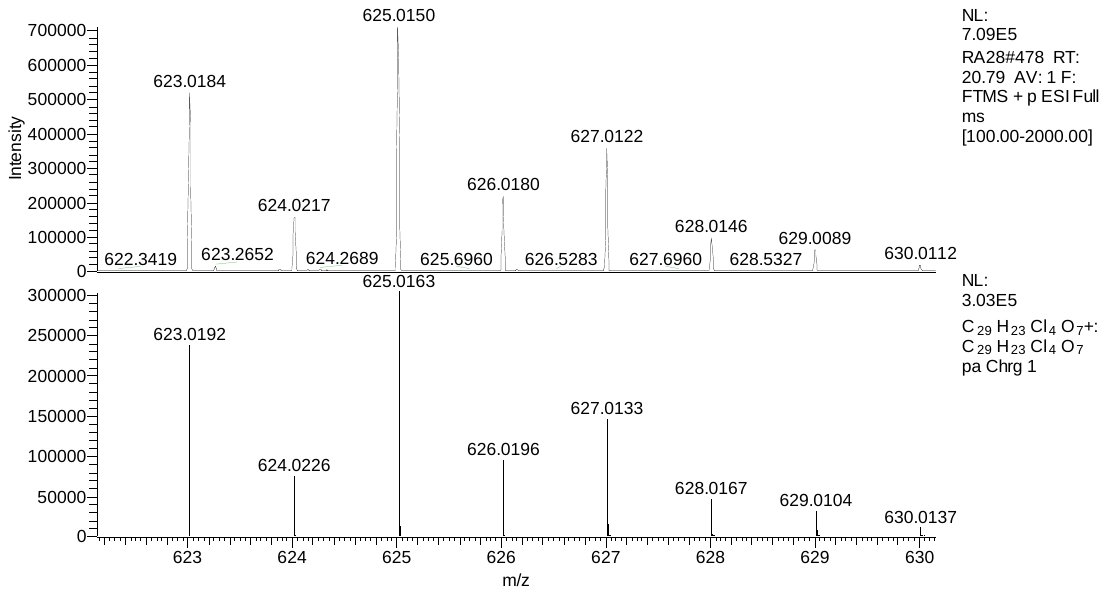

calculated

observed

Figure S70. LCMS Isotope Pattern of Accramycin K **11**

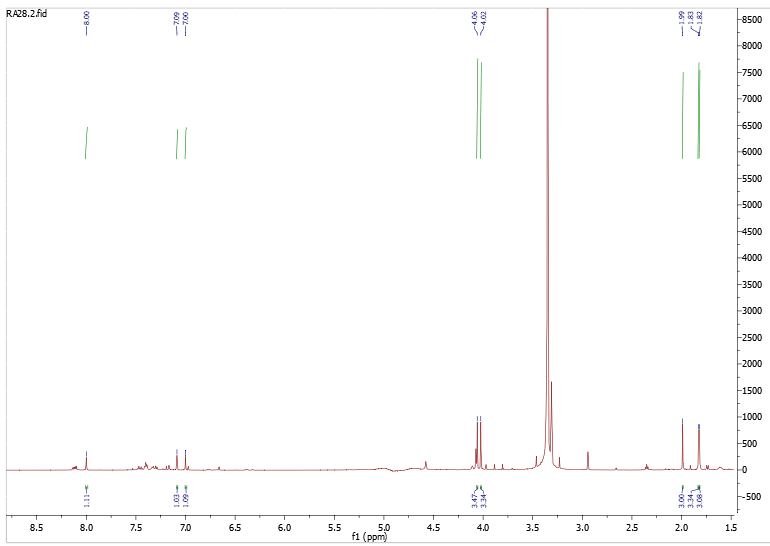

15

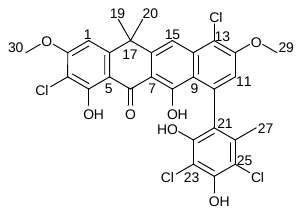

11

1

30

29

19, 20

27

Figure S71. ^1^H-NMR of Accramycin K **11** (CD_3_OD, 298K, 600MHz)

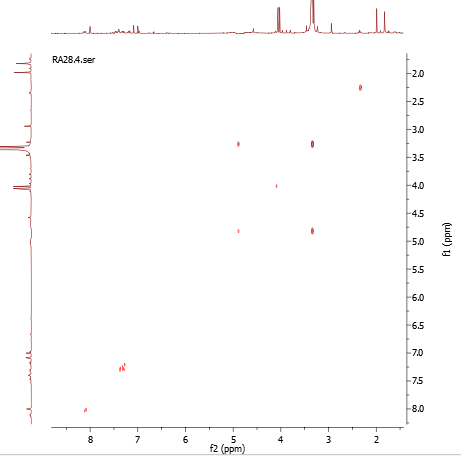

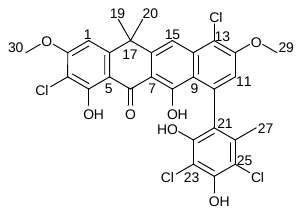

Figure S72. ^1^H-^1^H COSY of Accramycin K **11** (CD_3_OD, 298K, 600MHz)

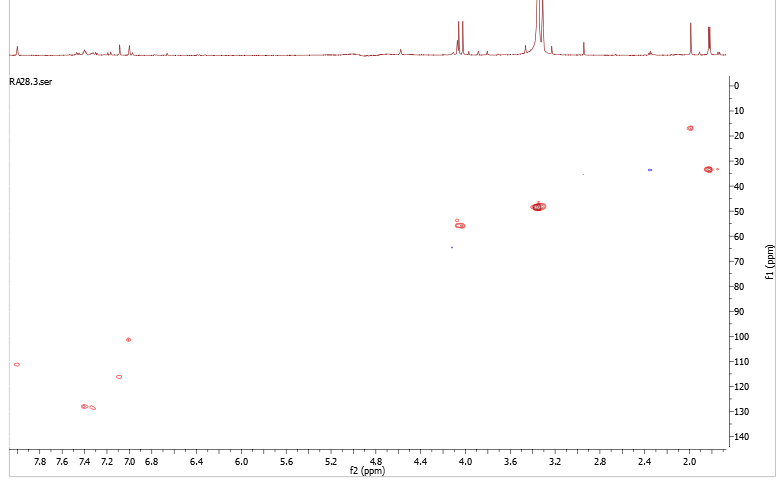

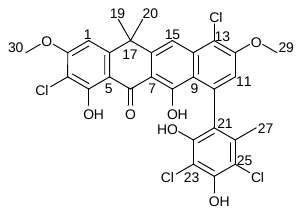

15

11

1

30

29

27

19, 20

Figure S73. HSQC of Accramycin K **11** (CD_3_OD, 298K, 600MHz)

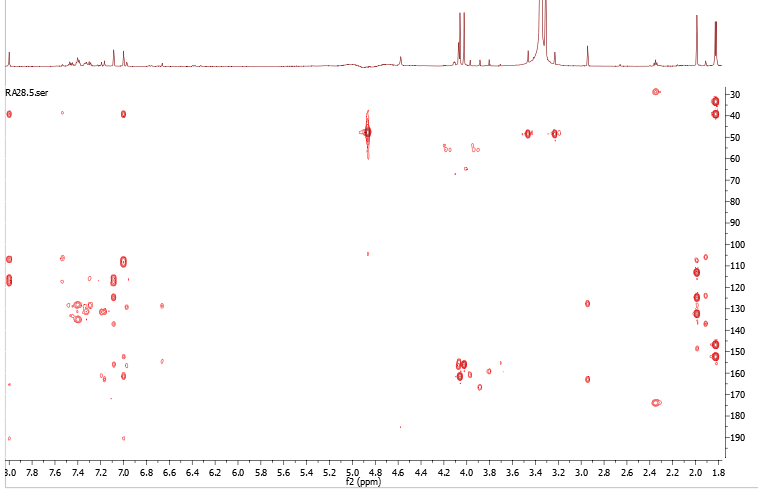

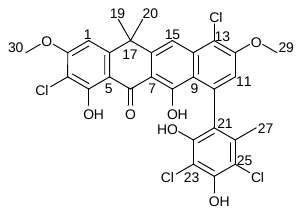

15

11

1

30

29

27

19, 20

Figure S74. HMBC of Accramycin K **11** (CD_3_OD, 298K, 600MHz)

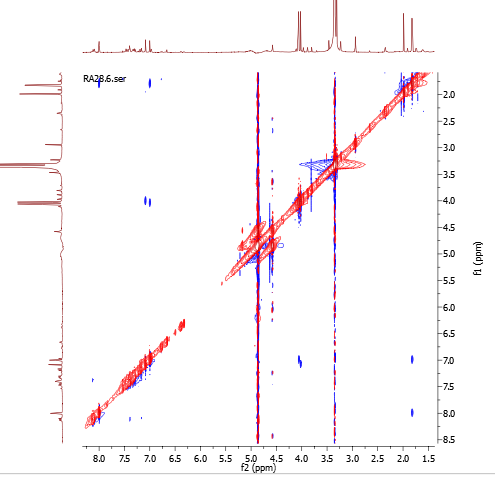

15

19, 20

27

30

29

11

1

11

1
